# Stemness overcomes IFNγ-induced antitumour exhaustion in human macrophages

**DOI:** 10.64898/2026.10.06.756629

**Authors:** Jessica Nitsche, Clara Jana-Lui Busch, Julien Subburayalu, Jeremy Favret, Anka Swiersy, Marie Hertrich, Fabian Rost, Daniela Ivan, Jiffin Benjamin, Mauro Franco, Qods Lahmar, Annemarie Scheede, Antonia Stammberger, Mariette Giannini, Yujun Zhang, Marc Schmitz, Ludwig Englmeier, Sethuraman Subramanian, Sandrine Sarrazin, Michael H. Sieweke

**Author notes:** Redpoint dx Germany, Erlenaustraße 11, 83080 Oberaudorf, Germany. These authors contributed equally and share first authorship.

## Abstract

Macrophages hold great promise for antitumour therapies^1–4^, particularly owing to their ability to infiltrate solid tumours^5^. Yet therapeutic success has been limited^6,7^, because macrophages exhibit rapid exhaustion of antitumorigenic activity in the tumour^5,8,9^. Here, we induced a stem-like self-renewal programme in human induced pluripotent stem cell (iPSC)-derived macrophages by deleting the self-renewal restricting transcription factors *MAFB* and *MAF* (MAF-DKO). Human MAF-DKO macrophages showed activated stem cell signatures, increased clonogenicity and proliferative potential. Increased self-renewal capacity was accompanied by enhanced mitochondrial fitness and invigorated energy metabolism. Importantly, these stemness features prevented exhaustion, cellular aging-associated parameters and loss of energy efficiency that are typically associated with proinflammatory macrophage activation. In addition, activated MAF-DKO macrophages showed a broad loss of IFNγ-triggered immunosuppressive pathways^10,11^, induced a super-activated state with robust cytotoxic activity and resisted exhaustion upon tumour cell exposure. Consequently, activated MAF-DKO macrophages showed intratumoral persistence with a stable antitumoral phenotype *in vivo,* conferred robust reduction of tumour burden and extended survival of tumourbearing mice. Together, this demonstrates that the induction of stemness in human macrophages can overcome critical roadblocks of antitumour therapy by conferring resistance to activation-triggered exhaustion of antitumour activity in the TME.

## Introduction

Macrophages represent an attractive target of cancer immunotherapy as they are abundant infiltrators of solid tumours and central regulators of the tumour microenvironment (TME)^5,8^. Their natural tumour-homing capacity has spurred particular interest in macrophage-based cellular therapies for solid tumours^1,2,4,6,12^, where therapeutic efficacy of engineered T cells has remained limited^13^.

Inflammatory activation has long been known to induce tumoricidal activity in macrophages^14,15^. A central inducer of this effect was found to be ‘macrophage activating factor’, later identified as IFNγ^16^. However, historic or recent efforts to harness this antitumoral potential^1–4^ have been clinically disappointing^6,7^, since activated macrophages rapidly lose antitumoral activity in the TME^5,8,9^. Much effort to overcome this problem has been focused on enhancing proinflammatory activation to reduce tumour-induced protumoral polarization^2,4,5,8,9^. However, IFNγ has not only antitumoral but also protumoral effects^17^ that trigger the acquisition of immunosuppressive pathways in recruited myeloid cells and tumour-associated macrophages (TAMs)^9–11,18^. Furthermore, IFNγ compromises proliferative capacity^19^ and metabolic stamina, as activated macrophages shift from oxidative phosphorylation (OX-PHOS) to glycolysis^20^. Given that glycolysis provides a much lower yield of ATP and is restricted to glucose as a nutrient source^21^, this shift dramatically erodes energy efficiency and metabolic flexibility in a highly nutrient-deprived TME^22–24^. Together, these phenomena rapidly exhaust persistence of macrophage antitumoral activity in the TME. Efforts to enhance inflammatory activation are therefore faced with the dilemma that IFNγ intrinsically triggers concomitant immunosuppressive pathways and antitumoral exhaustion in macrophages. This is of significant clinical interest as immune exhaustion has severely compromised the outcome of both historic and recent clinical trials with *ex vivo* IFNγ-stimulated^6^ or chimeric antigen receptor (CAR)-activated macrophages^7^.

Our previous work in mice showed that deletion of the self-renewal restricting transcription factors *MAFB* and *MAF* unlocks extended self-renewal capacity in differentiated macrophages^25,26^. Hypothesizing that activation of this stemness feature in macrophages could overcome the limitations of activation-driven exhaustion, we engineered human induced pluripotent stem cell (iPSC)-derived macrophages deficient for *MAFB* and *MAF* (MAF-DKO macrophages). Here, we show that self-renewal could also be activated in human macrophages by *MAFB* and *MAF* deletion. MAF-DKO macrophages exhibited gene expression signatures of self-renewing stem cells, showed enhanced clonogenicity, and unlocked proliferative potential. Strikingly, self-renewing MAF-DKO macrophages also activated additional features of stemness, including stress resistance, cytoprotective pathways, mitochondrial biosynthesis and activity. As a result, MAF-DKO macrophages resisted the typical sharp de-crease of OXPHOS experienced by conventional macrophages upon proinflammatory activation^20,27,28^, but rather responded with increased mitochondrial respiration and energy generation. The activation of stemness features also prevented cellular aging mechanisms, functional exhaustion, and immunosuppressive reprogramming of IFNγ-activated and tumour cell exposed MAF-DKO macrophages. IFNγ stimulation rather induced a super-activated state and cytotoxic programme that was effective against several tumour cell types. Consequently, MAF-DKO macrophages showed sustained tumoricidal activity *in vitro*, prolonged activation and persistence in tumours *in vivo* and mediated a potent restriction of tumour growth with extended survival of tumour-bearing hosts.

Our findings reveal a unique metabolic state of self-renewing human macrophages that respond to IFNγ activation with increased metabolic efficiency and mitochondrial respiration, resistance to functional exhaustion, and enhanced antitumour potency. Accordingly, MAF-DKO macrophages reunite distinct desirable functional features of antitumoral macrophages that were previously considered incompatible. Consequently, the activation of stemness properties overcomes a central restriction of macrophage antitumour activity and provides a promising strategy to stabilize and enhance the efficacy of macrophage-based therapies against solid tumours.

## Results

### Self-renewing macrophages show invigorated cellular fitness

To engineer self-renewal activity into human macrophages, we deleted the complete coding regions of the self-renewal restricting transcription factors *MAF* and *MAFB* in human iPSCs (Extended Data Fig. 1a-c). Using an established two-step embryoid body (EB)-based protocol^29,30^, we differentiated iPSCs into macrophages and maintained them in GM-CSF/M-CSF-containing culture medium (Extended Data Fig. 1d).

In order to characterize these cells, we performed RNAseq and gene set network analyses, which identified four major clusters of MAF-DKO macrophage-enriched terms (Fig. 1a). Consistent with our previous observations in mice^25,26^, human MAF-DKO macrophages were highly proliferative, as reflected in a strong enrichment of cell cycle terms and a major increase of cycling cells as measured by EdU-incorporation and PI staining (Fig. 1b-d; Extended Data Fig. 1e). Human MAF-DKO macrophages also showed a strong enrichment of self-renewing stem cell signatures (Fig. 1e) as well as self-renewal transcription factor regulons (Fig. 1f), including a core network centred around *MYC*, *KLF2*, and *KLF4* as we previously observed in mice^26^. Furthermore, single cells of MAF-DKO but not wild-type (WT) macrophages could give rise to clonally expanding colonies (Fig. 1g; Extended Data Fig. 1f). Together this activation of proliferative capacity, stem cell signatures, self-renewal regulons and clonal expansion potential indicated that human MAF-DKO macrophages had an analogous self-renewal potential as the equivalent mouse cells^25,26^.

**Figure 1:**
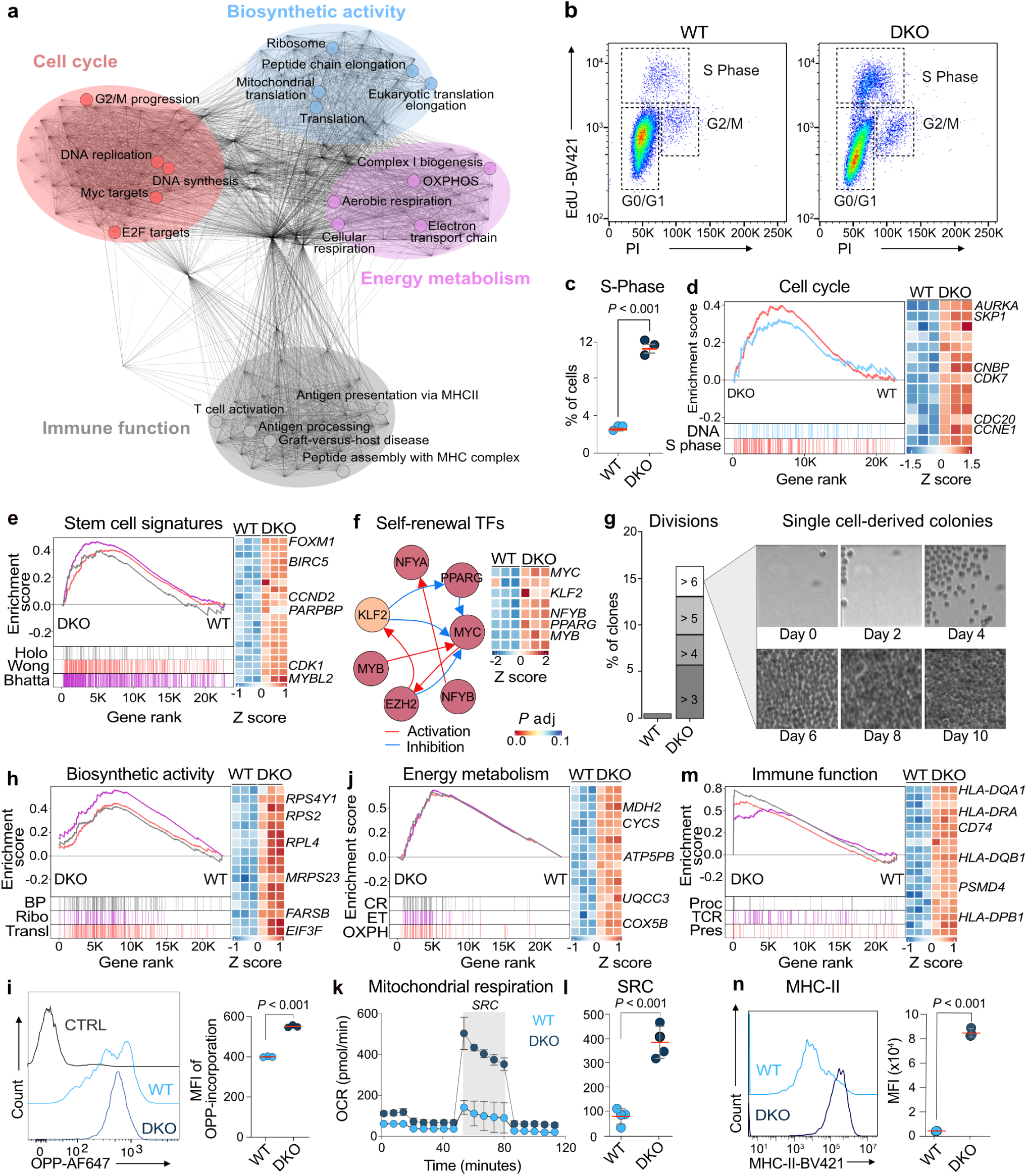
Activation of self-renewal in human MAF-DKO macrophages is linked to additional stemness features. **a,** RNAseq network analysis showing top 200 gene-sets enriched in MAF-DKO (*n* = 3) versus WT macrophages (*n* = 3) differentiated from iPSC (Extended Data Fig. 1d), Louvain community clustering, manually annotated by biological function (FDR < 0.05). **b,c** Representative FACS plot (**b**) and quantification (**c**) of percentages of CD45^HI^CD11b^HI^-macrophages in S-phase by EdU-incorporation (*n* = 3). **d,** GSEA of selected cell cycle gene sets from (**a**) and heatmap with Z-score-normalized expression of selected leading-edge genes. **e,** GSEA and leading-edge gene heatmap of selected stem cell modules (HOLO^106^, Wong^107^, Bhatta^108^). **f,** Network of self-renewal TFs activated in MAF-DKO macrophages shown in score normalized heatmap. Colour of circles indicates *P* value comparing activity in DKO vs WT with WT using t-test corrected with Benjamini-Hochberg. Lines indicate inferred decoupler-based crossregulation (activation in red, repression in blue). **g,** Single cell culture of EB harvest derived WT and MAF-DKO macrophages showing percentages of clones with indicated cell divisions at day 8, and example phase contrast images of MAF-DKO macrophages at indicated time points. **h,j,m**, Selected GSEA and heatmaps with Z-score-normalized expression of leadingedge genes as described in **d** for **h** biosynthetic activity: BP, biosynthetic process; Ribo, ribosome synthesis; Transl, translation. **j** energy metabolism: CR, cellular respiration; ET, electron transport chain; OXPH, oxidative phosphorylation. **m** immune function: Proc, antigen processing; TCR, TCR-signalling molecules; Pres, antigen presentation. **i**, Representative histograms and quantification of OPP-incorporation in WT or MAF-DKO macrophages (*n* = 3). **k,l,** Seahorse analysis of mitochondrial respiration in WT and MAF-DKO macrophages with quantification of SRC shown in grey (*n* = 4). **n,** FACS example and quantification of mean fluorescent intensity (MFI) for MHC-II expression in WT and MAF-DKO macrophages (*n* = 3). Statistical analysis was performed using unpaired two-tailed *t*-tests (**c**, **l**, and **n**). Data shown as mean ± s.d. and representative of at least three independent experiments, (**b,c,g,k,l,n)**. Full gene sets description shown in Extended Data Table 4. GSEA, gene set enrichment analysis. TF, transcription factor. EdU, 5-ethynyl-2’-deoxyuridine. OPP, O-propargyl-puromycin. SRC, spare respiratory capacity.

Surprisingly, however, human MAF-DKO macrophages showed additional, unexpected properties and gene expression changes that suggested further features of stemness beyond self-renewal capacity. These encompassed terms associated with enhanced biosynthetic activity, energy metabolism, and immune function (Fig. 1a,h-n). In line with enriched gene sets for ribosomal and protein synthesis, MAF-DKO macrophages showed a higher incorporation of the puromycin analogue O-propargyl-puromycin (OPP), indicative of enhanced capacity for mRNA translation (Fig. 1h,i). Furthermore, we found enrichment of terms for mitochondrial biogenesis, cellular respiration and OXPHOS (Fig. 1a,j), reflected in higher oxygen consumption rates (OCR) and ATP production detected by Seahorse metabolic assays (Fig. 1k; Extended Data Fig. 1g-j). Most strikingly, however, beyond the modestly increased basal respiration, MAF-DKO macrophages showed a dramatically increased spare respiratory capacity (SRC; Fig. 1k,l), indicating major reservoirs of OXPHOS and the potential to respond to increased energy demands that has been associated with stemness in other cell systems^31–33^. Finally, we identified increased terms for immune activation such as antigen pres-entation (Fig. 1a,m), which coincided with increased pan-MHC class II and HLA-DR expression levels on the cell surface of MAF-DKO macrophages (Fig. 1n; Extended Data Fig. 1k).

Together, these findings demonstrate that *MAF* and *MAFB* deficiency not only equipped human macrophages with self-renewal capacity but also with an increased biosynthetic activity, extensive reservoirs of energy production, and immune activation. Overall, this indicates a broad activation of stemness features associated with a surprising invigoration of general cellular fitness and immune cell functionality in these human macrophages.

### Stemness-activated macrophages resist IFNγ-induced exhaustion

We next investigated whether the enhanced cellular fitness of MAF-DKO macrophages impacted on the immune exhaustion experienced by IFNγ-activated macrophages in the TME. To address this, we performed scRNA sequencing of WT and MAF-DKO macrophages before and after IFNγ stimulation, followed by subsequent exposure to tumour cell-conditioned medium (TCM) as a surrogate of the immunosuppressive TME that activated macrophages encounter upon recruitment to the tumour. UMAP projections of the data revealed clearly distinct cellular states under the three conditions and between the two genotypes (Fig. 2a). Although both WT and MAF-DKO macrophages responded strongly to IFNγ, including canonical programmes of IFNγ or IFNγ/LPS-stimulated human blood monocytes (Extended Data Fig. 2a)^34,35^, they expressed unique and substantially different gene sets in response to IFNγ that were largely maintained after exposure to TCM (Fig. 2b-d; Extended Data Fig. 2b, Extended Data Table 1).

**Figure 2:**
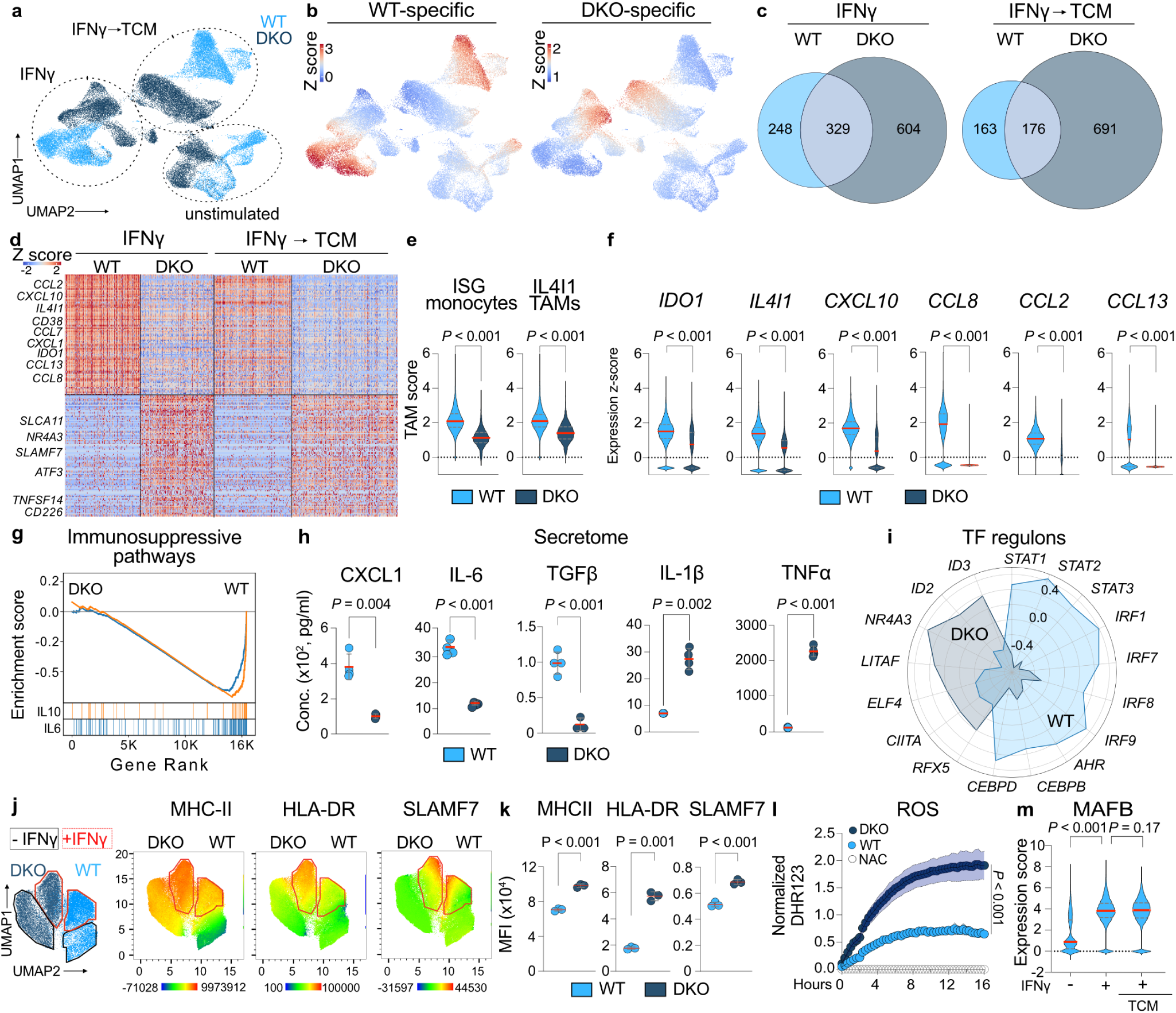
MAF DKO macrophages show a unique response to IFNγ. **a,b** UMAP projections of scRNAseq showing genotype and treatment conditions (**a**) and IFNγ-induced genes (**b**) in WT and MAF-DKO macrophages before and after stimulation with IFNγ followed by TCM. **c,** proportional Venn diagrams showing differential IFNγ-induced genes of WT and MAF-DKO macrophages in response to IFNγ and TCM. **d,** Heatmap of top100 IFNγ-induced DEGs for each genotype. **e,** Violin plots showing gene signature scores for ISG monocytes (ISG monos) and IL4I1^+^ TAMs (IL4I1 TAMs)^10^ in WT and MAF-DKO macrophages. **f,** Violin plots showing z-score expression of selected genes from (**d**). **g,** GSEA of *IL6* and *IL10* pathway gene sets in IFNγ-stimulated WT and MAF-DKO macrophages. **h,** Quantification of cytokine concentrations in supernatants of IFNγ-stimulated WT and MAF-DKO macrophages, *n* = 3 per group. **i,** Radar plot showing z-score of selected TF regulon activity^41^ in WT and MAF-DKO macrophages after IFNγ. **j,k,** UMAP projections and quantifications of selected surface marker expression in WT and MAF-DKO macrophages by spectral flow cytometry, *n* = 3 per group. **l,** Quantification of intracellular ROS levels in IFNγ-stimulated WT and MAF-DKO macrophages, monitored by DHR fluorescence signal in Incucyte SX5 live-cell imaging, expressed as relative green fluorescence per well normalised to cell confluence, *n* = 3 per group. **m,** Violin plots showing *MAFB* expression in WT macrophages in indicated conditions. Statistical analysis was performed using an unpaired twotailed *t*-tests (**e,f,h,k**), mixed-effect analysis (RELM model) (**l**) or two-way ANOVA with Tukey’s post hoc *t* test (**m**). Data shown as mean ± s.d. and representative of at least three independent experiments (**e,f,h,j,k,l,m**). Full gene set descriptions shown in Extended Data Table 4. TF, transcription factor. TCM, tumour cell conditioned medium. DHR, dihydrorhodamine 123. ROS, reactive oxygen species.

As a central proinflammatory cytokine, IFNγ is a key mediator of the antitumour response^17^ and is long known to activate tumoricidal activity in macrophages^3,14^. Yet, diverse efforts to harness this activity for antitumour therapy have been clinically disappointing^6,17^, possibly due to the intrinsically double-sided nature of IFNγ in tumour immunity, which includes both potent protumoral and immunosuppressive functions^11,17,18^. In this context, recent studies have identified monocyte-derived pro-inflammatory and immunosuppressive TAM populations associated with poor prognosis in many cancers^36–38^, in particular IFNγ-induced immunosuppressive IL4I1^+^ macrophages that develop from recruited monocytes with an interferon-stimulated gene (ISG) signature and orchestrate an immunosuppressive microenvironment in many tumours^10,39,40^. Our dataset revealed distinct differences between WT and MAF-DKO macrophages in such IFNγ responses. Whereas IFNγ-stimulated WT macrophages showed enriched expression of the previously characterized typical immunosuppressive signatures found in ISG monocytes and IL4I1^+^ TAMs (Fig. 2e, Extended Data Fig. 2c)^10^, including the critical immunosuppressive metabolic immune checkpoint genes *IDO1* and *IL4I1*, as well as the chemokines *CXCL9*, *CXCL10*, *CXCL11* (Fig. 2f; Extended Data Fig. 2d,e)^10,40^, MAF-DKO macrophages were resistant to this IFNγ response. Remarkably, they also failed to induce critical mediators of myeloid-derived suppressor cells (MDSCs)^1,2,11,18^ including *CXCL1*, *CXCL2*, *CXCL5*, *IL6*, and *IL10* (Fig. 2f; Extended Data Fig. 2e), or mediators of protumoral, monocyte-derived TAMs^1,2^, including *CCL2*, *CCL7*, *CCL8*, and *CCL13* (Fig. 2f; Extended Data Fig. 2d,e; Extended Data Table 1). Consistent with this, IFNγ-stimulated MAF-DKO macrophages also did not induce immunosuppressive pathways, including *IL6* and *IL10* signalling (Fig. 2g, Extended Data Fig. 2f) and secreted lower amounts of CXCL1, IL-6, and TGFβ (Fig. 2h). These observations were further supported by decoupler analysis which predicts cell-state determining transcription factors (TFs)^41^. TF regulons known to mediate the IFNγ-induced immunosuppressive programme of MDSCs (*STAT1*, *STAT3*, *CEBPB* and *CEBPD*), ISG monocytes and IL4I1^+^ TAMs (*STAT1*, *STAT2*, *IRF1*, *IRF7*, *IRF8*, *IRF9*, and *AHR*) were activated in IFNγ-stimulated WT macrophages but not induced in MAF-DKO macrophages (Fig. 2i)^2,10,37,42–45^. *STAT1* and the aryl hydrocarbon receptor (*AHR*) have been shown to induce IL4I1, which in turn generates ligands for AHR from L-tryptophan^38^. Importantly, macrophage AHR activity suppresses antitumour immunity, enhances tumour aggressiveness^38,42^, and is associated with a poor prognosis in many tumours^38^. Together, this indicated that MAF-DKO macrophages were broadly resistant to a wide spectrum of IFNγ-induced immunosuppressive pathways associated with tumour progression, including the critical IL4I1/AHR-axis.

Despite this remarkable resistance of MAF-DKO macrophages to IFNγ-induced immunosuppressive pathways, they activated multiple inflammatory and immune function pathways after IFNγ stimulation (Extended Data Fig. 2f). Decoupler analysis^41^ also identified a unique network of immune function TF regulons specifically in IFNγ-stimulated MAF-DKO macrophages (Fig. 2i), such as antitumoral macrophage activity (*ID3*)^46^, proinflammatory macrophage activation (*ID2*, *NR4A3*/*NOR1*)^47,48^, IFNγ-dependent inflammatory amplification (*LITAF*, *ELF4*)^49,50^, and antigen presentation (*CIITA*, *RFX5*)^51^. Consistent with this, we observed increased surface marker expression of proinflammatory activation, such as MHC class II molecules, including HLA-DR, and SLAMF7 (Fig. 2j,k), an effector molecule of superactivation in macrophages that triggers an inflammatory signalling cascade culminating in enhanced TNFα production^52^. In line with this, we found dramatically increased secretion of proinflammatory cytokines, in particular over 20-fold higher levels of TNFα (Fig. 2h). Further reflecting a proinflammatory activation^20^, MAF-DKO macrophages also showed strongly increased ROS production in response to IFNγ or PMA (Fig. 2l; Extended Data Fig. 2g), whereas phagocytosis was unaffected (Extended Data Fig. 2h).

Further probing the underlying mechanism of this unique IFNγ response, we investigated *MAF* and *MAFB* expression in WT macrophages. Similar to previous reports^35^, we observed that in WT macrophages *MAF* expression and *MAF* target genes were transiently repressed by IFNγ stimulation and reactivated after exposure to TCM (Extended Data Fig. 2i-k). By contrast, *MAFB* expression was increased by IFNγ stimulation in WT macrophages (Fig. 2m), paralleling the described immunosuppressive ISG monocyte and IL4I1^+^ TAM programmes (Fig. 2e, Extended Data Fig. 2c). This demonstrates a critical difference of the two TFs and suggests that loss of *MAFB* specifically prevents the activation of IFNγ-dependent immunosuppressive programmes that are insensitive to the loss of *MAF*.

Together our data show that the combined absence of *MAF* and *MAFB* in macrophages comprehensively abolished IFNγ-induced immunosuppressive programmes, while simultaneously enabling pathways of proinflammatory super-activation, thus effectively uncoupling immunosuppressive and proinflammatory IFNγ effects in macrophages.

### Stemness resolves metabolic macrophage activation dilemma

OXPHOS is generally suppressed by proinflammatory activation^20,53^, as IFNγ-stimulated macrophages redirect their metabolism from OXPHOS to glycolysis, sacrificing efficient energy provision for rapid proinflammatory effector functions^53,54^. We therefore investigated energy metabolism in IFNγ-stimulated MAF-DKO or WT macrophages by assessing OCR (Fig. 3a; Extended Data Fig. 3a) and extracellular acidification rates (ECAR; Extended Data Fig. 3b), indicative of OXPHOS and glycolytic metabolism, respectively. Surprisingly, rather than decreasing energy efficiency, MAF-DKO macrophages further increased their energy metabolism in response to IFNγ, showing massive increases in basal and maximum respiration rates (Fig. 3a,b; Extended Data Fig. 3a), as well as enhanced ECAR (Extended Data Fig. 3b). As a result, MAF-DKO macrophages massively increased their ATP production by virtue of both enhanced glycolysis and OXPHOS (Fig. 3c), combining an increase in both rapid (glycolysis) and efficient energy provision (OXPHOS). Importantly, MAF-DKO macrophages also maintained a strongly increased ECAR, OCR, basal and maximal respiration, as well as ATP production after subsequent exposure to TCM (Extended Data Fig. 3c-g).

**Figure 3:**
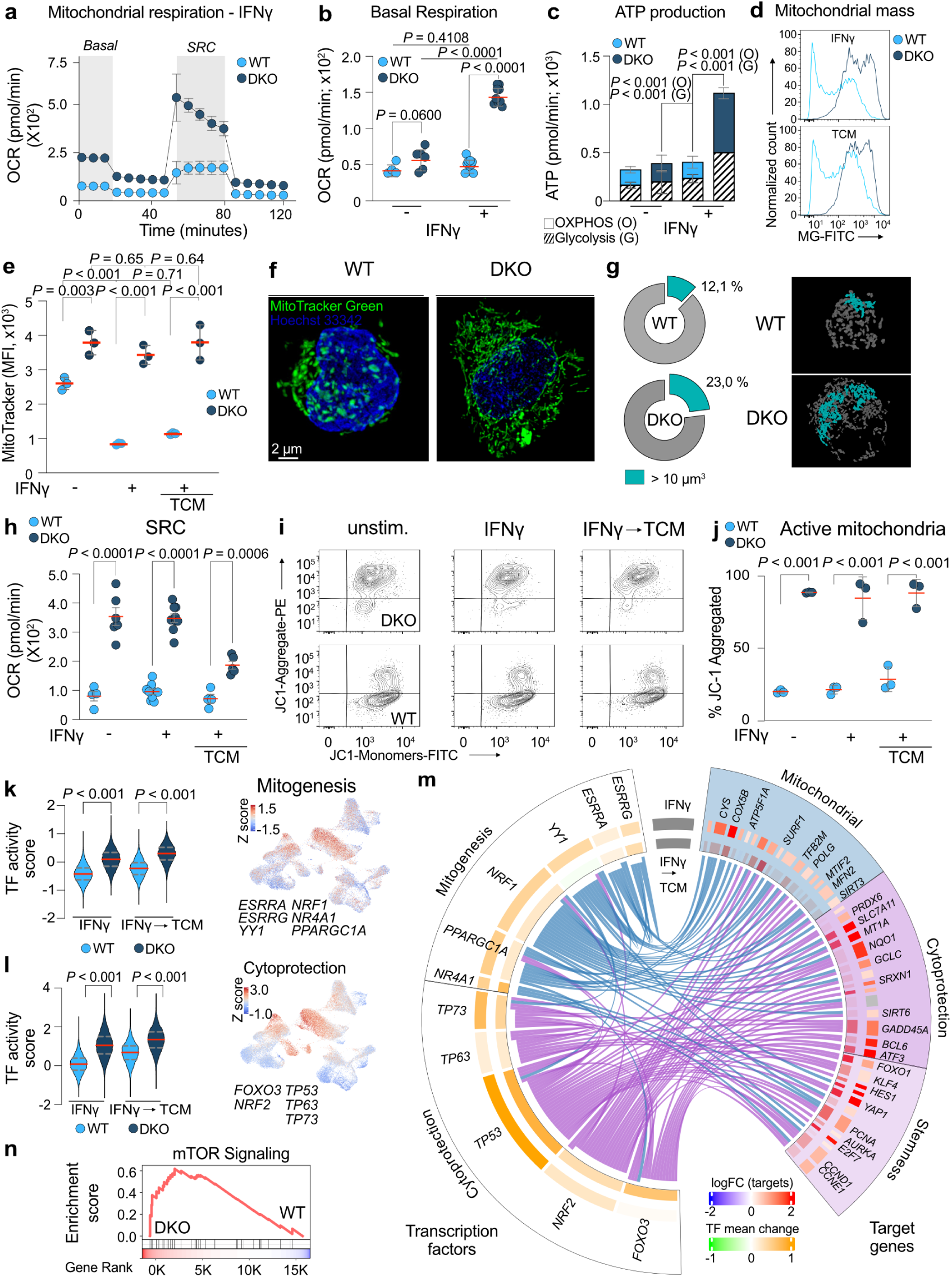
Stemness overcomes metabolic macrophage activation dilemma. **a,** Seahorse analysis of mitochondrial respiration in WT and MAF-DKO macrophages 16 hours after IFNγ stimulation with basal (basal) and spare respiration (SRC) highlighted in grey, *n* = 4-6 per group. **b,** Quantification of basal respiration, *n* = 4-6 per group. **c,** Quantification of glycolytic and mitochondrial ATP production in WT and MAF-DKO macrophages, *n* = 4-6 per group. **d,e,** Representative histograms and quantification of MitoTracker Green FM staining, reflecting mitochondrial mass under indicated conditions in WT and MAF-DKO macrophages, *n* = 3 per group. **f,g,** Representative confocal microscopy images of Mito-Tracker Green FM-stained WT or MAF-DKO macrophages, quantification of large mitochondrial networks and representative mitochondrial segmentation images after Nellie segmentation^109^. **h,** Quantification of SRC under indicated conditions in WT and MAF-DKO macrophages. **i,j,** Representative contour plots (**i**) and percentage (**j**) of live macrophages positive for JC-1 aggregates in WT and MAF-DKO macrophages, *n* = 3 per group. **k,l,** Violin plots and UMAP projections of indicated TF activity scores in WT and MAF-DKO macrophages associated with mitogenesis and cytoprotection. **m,** CIRCOS plot showing TF activity and differential gene expression (log_2_FC) in associated positive target genes in MAF-DKO macrophages over WT macrophages after exposure to IFNγ or subsequent exposure to TCM. **n,** GSEA mTOR signalling comparing MAF-DKO IFNγ and TCM vs. IFNγ and TCM in WT macrophages. Statistical analysis was performed using unpaired two-way ANOVA with Tukey’s post hoc *t* test (**b,c,e,h,j,k,l**). Data shown as mean ± s.d. and representative of at least three independent experiments (**a,b,c,d,e,h,I,j**). Full gene set descriptions shown in Extended Data Table 4. OCR, oxygen consumption rate. SRC, spare respiratory capacity. TF, transcription factor.

Thus, MAF-DKO macrophages appear to engage their strong reservoir of mitochondrial respiratory capacity for a highly unusual increase in energy production following proinflammatory activation with IFNγ and subsequent exposure to TCM.

### Macrophage stemness associated robust mitochondrial fitness

Mitochondrial dysfunction drives glycolytic reprogramming and functional exhaustion in T cells^55,56^. Reversely, mitochondrial transfer can confer stem-cell-like self-renewal characteristics, as well as energy and antitumoral efficiency^57^. Furthermore, selection of T cells for healthy mitochondria can identify cells with enhanced stemness^31^ and activation of stemness is associated with increased mitochondrial health^33^. Given the apparent role of mitochondrial fitness as a feature of stemness^31,58^, and the remarkably robust energy metabolism of MAF-DKO macrophages, we investigated mitochondrial biology in response to macrophage activation and tumour cell exposure. Using MitoTracker Green FACS staining, we observed that MAF-DKO macrophages had significantly higher mitochondrial mass under all conditions and, importantly, were resistant to the major decline seen after IFNγ stimulation and TCM exposure in WT cells (Fig. 3d,e). By investigating the mitochondrial morphology of IFNγ-stimulated macrophages, we observed an intricate network of elongated, frequently fused mitochondria in MAF-DKO macrophages, whereas in most WT cells mitochondria had a fragmented, rounded appearance, also reflected in the smaller size and reduced number of large mitochondrial networks (Fig. 3f,g). Mitochondrial fragmentation is an indicator of poor adaptation capacity to stress conditions^32,59^. To quantitatively assess this, we compared SRC, which is a particularly robust functional parameter of mitochondrial reserve that correlates with mitochondrial fragmentation^32^. Indeed, SRC was strongly increased under all conditions in MAF-DKO macrophages, including after IFNγ activation and TCM exposure (Fig. 3h). Furthermore, the fluorescent dye JC-1, an indicator of mitochondrial membrane potential (MMP), revealed a very high proportion (> 85%) of JC-1 Red^+^ cells, indicative of active mitochondria with intact electron transport chain and ATP-producing capacity in MAF-DKO macrophages (Fig. 3i,j). This was a vast increase compared to the small proportion of cells with active mitochondria (< 20%) in WT macrophages that harboured a majority of dysfunctional mitochondria indicative of metabolic stress and impaired mitochondrial respiration (Fig. 3i,j).

Together, these results suggest that the observed invigorated energy metabolism in MAF-DKO macrophages, even after proinflammatory activation with IFNγ and subsequent exposure to TCM, is enabled by enhanced mitochondrial integrity and function. Therefore, MAF-DKO macrophages appear to exhibit strong stemness features of mitochondrial fitness that render macrophage activation compatible with a robust energy metabolism and resistant to activation-induced exhaustion in the TME.

### Stemness-induced mitogenesis and cellular stress resistance

In order to unravel the molecular basis for the mitochondrial fitness and resistance to metabolic exhaustion, we further analysed the genes that were specifically activated in MAF-DKO macrophages but absent in WT macrophages after IFNγ stimulation and subsequent exposure to TCM. Besides enriched terms for proliferation, self-renewal, biosynthetic activity, and immune function, IFNγ-activated MAF-DKO macrophages also showed terms for mitochon-drial respiration and function (Extended Data Fig. 2f). Interestingly, analysis by decoupleR^41^ revealed that IFNγ-stimulated and TCM-exposed MAF-DKO macrophages activated interlinked regulons associated with stem cell programs, mitochondrial biogenesis, cellular longevity, and stress protection programmes (Fig. 3k-m, Extended Data Fig. 3h).

Remarkably, upon IFNγ-stimulation and exposure to TCM, MAF-DKO macrophages activated a comprehensive mitochondrial biogenesis programme encompassing virtually every stage of mitochondrial generation, maintenance and function (Fig. 3k,m). The specifically activated genes identified central regulons of mitochondrial biology driven by the nuclear factors *ESRRA*, *ESRRG*, *YY1*, *NRF1*, *PPARGC1A* and *NR4A1* (*NUR77*)^55,60–64^. The target genes included key components of mitochondrial respiration, such as *NDUFS8* (complex I), *CYCS* (the electron carrier between complex III and complex IV), *COX5B* and *COX7C* (complex IV), and *ATP5F1A* (ATP synthase, complex V). Activated genes were also involved in mitochondrial protein import and respiratory chain assembly (*TOMM34*, *SURF1*), mitochondrial DNA replication and transcription (*POLG*, *TFAM*, *TFB2M*), mitochondrial ribosome biogenesis and translation (*TFB1M*, *MTIF2*), as well as mitochondrial fusion (*MFN2*) and mitochondrial stress resistance and longevity (*SIRT3*) (Fig. 3m). Furthermore, we also observed enriched terms of mTOR signalling (Fig. 3n), which is as a key upstream activating pathway of *PPARGC1A*, is a central TF driving *de novo* mitochondrial biogenesis and metabolism^55,62,64^, and controlling one of the major activated regulons observed in MAF-DKO macrophages (Fig. 3m).

IFNγ-stimulated and TCM-exposed MAF-DKO macrophages also engaged regulons driven by *FOXO3*, *NFE2L2/NRF2*, and *p53* family members (Fig. 3l,m), central regulators of longevity, oxidative stress resistance and genome protection, respectively^65–69^. The associated target genes revealed comprehensive and integrated oxidative and replicative stress resistance mechanisms encompassing antioxidant defence, redox homeostasis, DNA repair, and metabolic adaptation. The genes included longevity-associated sirtuins (*SIRT1*, *SIRT3* and *SIRT6*), stress-responsive TFs (*ATF3*, *FOXO1*, and *BCL6*), antioxidant enzymes (*GPX3*, *PRDX5*, *PRDX6*, *NQO1*, *SRXN1*), glutathione biosynthesis and recycling components (*SLC7A11*, *GCLC*, and *GSS*), mediators of DNA repair and cellular stress adaptation (*GADD45A*, *SESN3*, and *MT1A*), providing a molecular rationale for how MAF-DKO macrophages could tolerate high mitochondrial activity (Fig. 3a-c; Extended Data Fig. 3a,d-g), IFNγ-induced ROS (Fig. 2l), and proliferation-induced replication stress (Fig.1b,c).

Notably, we observed a significant cross-regulation between these regulons, resulting in the integration of mitogenesis and cytoprotection with proliferation (*PCNA*, *AURKA*, *E2F6*, *E2F7*, *CCND1*, and *CCNE1*), and stemness pathways (*FOXO1*, *KLF4*, *YAP1*, *HES1*, *FBXW7*). This was in line with the activation of regulons of TFs associated with cellular longevity of stem cells (Extended Data Fig. 3h)^68,70–74^, including HES transcription factors that have been associated with p53-mediated cytoprotection in stem cells^65^.

Collectively, this suggests that MAF-DKO macrophages integrate mitochondrial biogenesis, cellular longevity and stress resistance pathways as comprehensive features of stemness activation. Together this provides a molecular rationale for the observed mitochondrial fitness and resistance to the metabolic exhaustion typically experienced by IFNγ-activated macrophages in the TME.

### Stemness-activated macrophages resist aging and exhaustion

Proinflammatory activation of macrophages can engage similar gene expression programmes as aging^75^, including IFN-I signatures^76^, senescence-associated secretory phenotypes (SASP) and upregulation of CD38, an aging-associated nicotinamide adenine dinucleotide (NAD^+^)-consuming enzyme^10,77,78^. Furthermore, senescence of TAM populations has been shown to promote tumorigenesis in lung cancer models^79,80^. Interestingly, the response induced by CAR vectors in human macrophages shares similarities with IFNγ-activated IFN-I response signatures^12,76,81^, which might contribute to the sensitivity of human CAR-transduced macrophages to exhaustion^7^. We therefore analysed IFNγ-driven macrophage activation-induced cellular aging and exhaustion pathways. Indeed, IFNγ-activated WT macrophages showed an induction of both canonical and macrophage-specific senescence signatures, IFN-I and CAR-induced responses, which were collectively absent in IFNγ-activated MAF-DKO macrophages (Fig. 4a-c; Extended Data Fig. 4a-f). This resistance to senescence and macrophage-specific aging signatures^75,79,80,82^ included CD38, both on the message and protein expression level (Fig. 4c-e). Consistent with the reduced levels of this NADase, IFNγ-activated MAF-DKO macrophages showed higher levels of intracellular NAD^+^ than WT cells (Fig. 4f). Analysis by decoupler^41^ further revealed that IFNγ-stimulated WT macrophages activated regulons associated with an IFN-I response (*STAT1, STAT2, STAT3, IRF3*, *IRF7*, *IRF8*, and *IRF9*)^76,83^, CD38 activation (*STAT1, STAT2, STAT3*)^84^, metabolic exhaustion (*HIF1A*)^20^, SASP and aging signatures (*AHR*, *CEBPD* and *CEBPB*)^85^ (Fig. 4g). Furthermore, upon exposure to TCM, WT macrophages shifted to a different set of exhaustion regulons including the self-renewal restricting factors *BATF*^86,87^, *MAF* and *MAFB*^25,26^, and *LXR*/*RXR* factors (*NR1H3* (*LXR*α), *NR1H2* (*LXR*β), *NR1H4*, *RXRA, RXRB*), central activators of CD38 in macrophages^84,88^ (Fig. 4h). Remarkably, MAF-DKO macrophages were resistant to both of these IFNγ-activation- and TCM-induced exhaustion networks (Fig. 4g,h).

**Figure 4:**
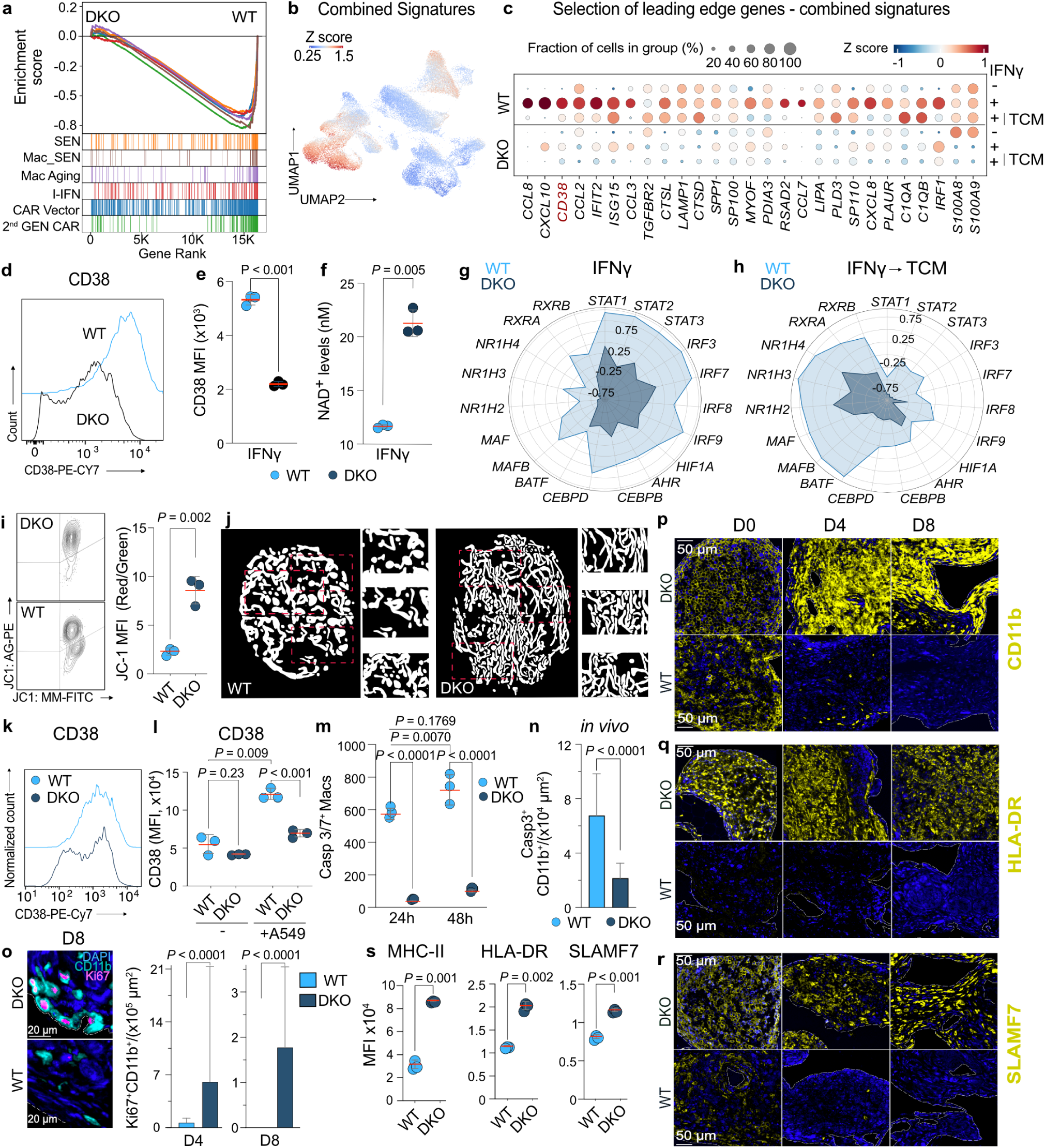
MAF-DKO macrophages resist exhaustion and activity loss in the TME. **a,b,** GSEA (**a**) and combined UMAP projection (**b**) of published signatures of senescence (SEN^82^, Mac_SEN^79^, Mac Aging^75^, IFN-I, CAR Vector^12^ and 2^nd^ GEN CAR^81^) under IFNγ and TCM conditions in WT and MAF-DKO macrophages. **c,** Bubble plot indicating the fraction of cells of selected leading-edge genes from combined gene signatures of (**a**). **d,e,** Representative histograms and MFI quantification of CD38 surface expression in WT and MAF-DKO macrophages 24 h after IFNγ stimulation. **f,** Colorimetric quantification of intracellular NAD^+^ levels 24 h after IFNγ stimulation. **g,h,** Radar plots showing selected exhaustion-related transcription factor activity after IFNγ stimulation (**g**) or subsequent TCM treatment (**h**). **i,** Representative contour plots and quantification of JC1-Red/Green WT and MAF-DKO macrophages after IFNγ activation and two days of subsequent co-culture with A549 cells, *n* = 3 per group. **j,** Representative mitochondrial segmentation images from WT and MAF-DKO macrophages, showing whole cells and inserts at higher magnification, based on live Mito-Tracker Green FM-staining of processed 3D confocal-images and segmented using MoDL^99^. **k,l,** Representative histograms and MFI quantification of CD38 surface expression in WT and MAF-DKO macrophages in monoculture or two days after co-culture with A549 cells at an E:T ratio of 10:1, *n* = 3 per group. **m,** Caspase 3/7^+^ macrophages after 24 hours of IFNγ-activation and further 24 hours or 48 hours of co-culture with A549 cells, expressed as counts per image using a Caspase 3/7 sensitive dye and SX5 Incucyte analyser, *n* = 3 per group. **n**, Casp3^+^CD11b^+^ WT or MAF-DKO macrophages per area 24 hours after IFNγ-activation and co-transplantation with A549 cells *in vivo*. **o,** Ki67^+^CD11b^+^ WT or MAF-DKO macrophages per area at indicated time points after IFNγ activation and co-transplantation with A549 cells *in vivo*. **p-r,** CD11b (**p**), HLA-DR (**q**), and SLAMF7 (**r**) of IFNγ-activated WT or MAF-DKO macrophages at indicated time points after co-transplantation with A549 cells *in vivo*, representative of two independent experiments, with at least two mice per genotype and timepoint per experiment. **s,** Flow cytometric MFI quantification of indicated cell surface markers after IFNγ stimulation and subsequent 96 hours exposure to TCM in WT and MAF-DKO macrophages, *n* = 3 per genotype. Statistical analysis by unpaired two-tailed t-tests (**e,f,j,n**,**o,s**), or a two-way ANOVA with Tukey’s post hoc t test (**l, m)**. Data shown as mean ± s.d. or median ± IQR **(n,o**). and representative of at least three independent experiments (**d,e,f,k,l,m,s**). Full gene set descriptions shown in Extended Data Table 4. TCM, tumour cell conditioned medium.

To address whether the concomitant activation of stemness-associated metabolic invigoration, mitochondrial fitness and resistance to cellular aging pathways could improve macrophage persistence in competition with tumour cells, we performed co-culture experiments of IFNγ-activated macrophages with A549 cells, a human NSCLC-derived cell line. Strikingly, in contrast to WT cells, MAF-DKO macrophages did not develop mitochondrial dysfunction as indicated by maintained MMP and lack of morphological fragmentation (Fig. 4i,j). They also showed strongly reduced levels of CD38 or Caspase 3/7 (Fig. 4k-m). Together this indicated an extraordinary mitochondrial robustness and resistance to exhaustion and cell death mechanisms in the presence of tumour cells. To determine whether this was also the case *in vivo*, we co-injected macrophages and A549 cells subcutaneously into the flank of NSG-SGM3 mice. Compared to WT macrophages, IFNγ-stimulated MAF-DKO macrophages also showed reduced numbers of Caspase 3^+^ cells *in vivo* (Fig. 4n) and increased numbers of Ki67^+^ cells, at least up to 8 days (Fig. 4o; Extended Data Fig. 5a,b), indicating reduced apoptosis and maintained proliferative capacity in the TME. Consequently, WT macrophages rapidly declined and were nearly completely depleted by 8 days post injection, whereas MAF-DKO macrophages showed a high persistence and were retained *in vivo* at strikingly increased numbers (Fig. 4p; Extended Data Fig. 4g, 5c). Since macrophages in the TME typically show loss of antitumorigenic function^8^, we investigated whether MAF-DKO macrophages also showed persistent proinflammatory activation in the presence of tumour cells. Importantly, MAF-DKO macrophages retained expression of HLA-DR and SLAMF7 *in vitro* and *in vivo* (Fig. 4q-s; Extended Data Fig 4h,i, 5d,e).

Together, these results indicate that resistance to multiple aging, cell death and exhaustion mechanisms enables IFNγ-stimulated MAF-DKO macrophages to persist in a hostile environment of tumour cells while retaining proliferative capacity and a proinflammatory, antitumoral phenotype.

### Stemness-activated macrophages exert tumoricidal activity

To test whether stemness associated energy efficiency, enhanced effector functions and cellular persistence conferred improved antitumour immunity to MAF-DKO macrophages, we analysed IFNγ-induced tumoricidal macrophage activity^3^. Using K562 tumour cells, a classical target for cytotoxicity assays, WT macrophages showed no cytotoxic activity without IFNγ, and only a delayed marginal effect 48 hours after IFNγ stimulation (Fig. 5a-c). By contrast, MAF-DKO macrophages already showed some cytotoxicity in the absence of IFNγ (Fig. 5a), with a massively increased cytotoxic effect after activation with IFNγ, resulting in a rapid and near total elimination of K562 cells in a time and dose-dependent manner within less than 20 hours (Fig. 5b,c; Extended Data Fig. 6a; Supplementary Video 1). Importantly, MAF-DKO macrophages demonstrated an antitumoral phenotype also against solid tumour cells, as exemplified by A549 NSCLC cells and SK-OV-3 ovarian carcinoma cells (Fig. 5d,e; Extended Data Fig. 6b,c). Moreover, MAF-DKO macrophages showed a durable antitumoral potential, since they could still induce apoptosis and secondary necrosis in A549 cells after 5 days of co-culture (Extended Data Fig. 6d). Together, these observations indicate that MAF-DKO macrophages are particularly sensitive to the induction of tumoricidal activity by IFNγ, enabling the activation of a robust and persistent antitumoral activity against multiple tumour cell types.

**Figure 5:**
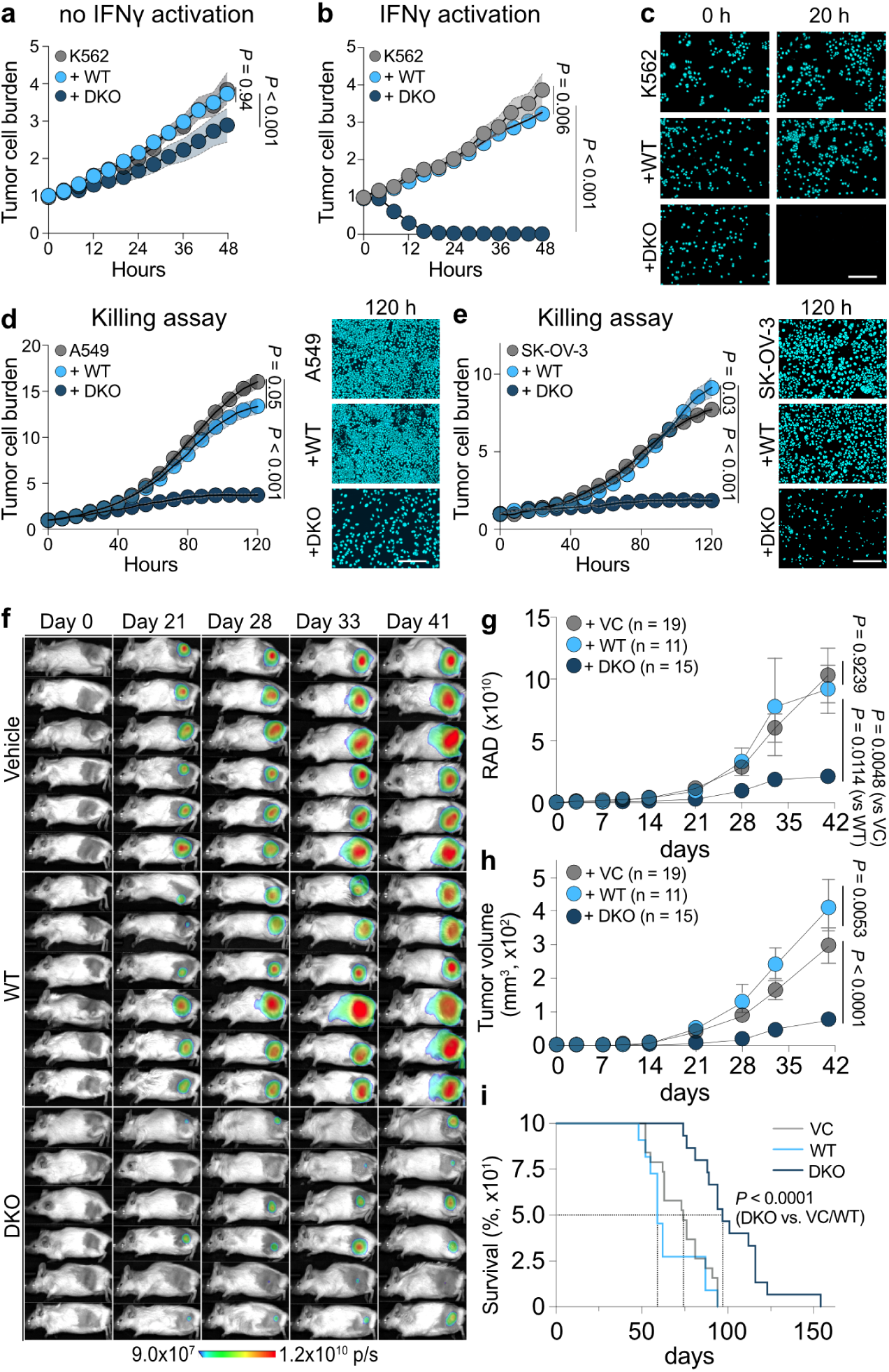
MAF-DKO macrophages exert robust antitumour activity *in vitro* and *in vivo*. **a-c,** Cytotoxicity assay quantifying NIR^+^ K562 tumour cell numbers over time using an SX5 Incucyte instrument. K562 cells were cultured without or with WT or MAF-DKO macrophages either IFNγ naïve (**a**) or after 16 hours of IFNγ stimulation (**b**) at an effector-to-target (E:T) ratio of 10:1. Values are normalised to time = 0 hours, *n* = 3 per group. **c,** Representative images of NIR^+^ tumour cells for indicated time points from (**b**). Scale bar = 20 µm. **d,e,** Incucyte-based assessment and representative images of NIR^+^ A549 (**d**) or SKOV-3 cells (**e**) cultured as outlined in (**b**). Values are normalised to time = 0 hours, *n* = 3 per group. Scale bar = 20 μm. **f,** Representative bioluminescent images (BLI) of mice transplanted with A549 cells and vehicle or IFNγ-stimulated WT or MAF-DKO macrophages after indicated time points with an E:T ratio of 50:1. **g,** Average BLI (flux p/s) of mice treated with vehicle (*n* = 19), WT (*n* = 11), or MAF-DKO macrophages (*n* = 15) over time. **h**, Non-invasive, calliper-based assessment of tumour growth in mm^3^ of mice shown in (**f**) as described previously^100^. **i,** Survival of vehicle (*n* = 19), WT (*n* = 11), or MAF-DKO macrophage transplanted mice based on reaching termination criteria for tumour-bearing mice analysed by log-rank Mantel-Cox test, *P* < 0.0001. Statistical analysis was performed using unpaired two-tailed t-tests or Mixed-effect analysis (RELM model) to compare two groups over time (**a**,**b**,**d**,**e**) or a two-way ANOVA with Tukey’s post hoc t-test (**g,h**). Data shown as mean ± s.d. (**a,b,d,e**) or median ± IQR (**g,h**). Representative results from one of at least two (**g,h**) or three independent experiments (**a-e**). NIR, near-infrared. VC, vehicle control.

To evaluate the *in vivo* relevance of these observations, we transplanted A549 tumour cells subcutaneously into NSG-SGM3 mice together with vehicle control, WT or MAF-DKO macrophages and followed progression of luciferase-transduced A549 cells by bioluminescent imaging (BLI) and calliper-based volumetric measurements (Extended Data Fig. 7a). Notably, mice treated with MAF-DKO macrophages exhibited a marked reduction in tumour burden as reflected in both decreased BLI signal intensity and smaller tumour volumes (Fig. 5f-h; Extended Data Fig. 7b,c). Consistently, even such a single co-injection of MAF-DKO macrophages with A549 NSCLC cells resulted in a significantly prolonged overall survival of up to 50 days in tumour-bearing mice (Fig. 5i). Further analysis indicated that these antitumoral effects of MAF-DKO macrophages were similar in both sexes, demonstrating that MAF-DKO macrophages conveyed a durable, sex-independent suppression of tumour growth *in vivo* (Extended Data Fig. 7d-i).

Together, these data show that activation of stemness in human MAF-DKO macrophages was associated with a broad antitumoral phenotype *in vitro* and *in vivo*.

## Discussion

Here, we report for human macrophages that the activation of stemness features with enhanced mitochondrial fitness unlocks a stable antitumour activity resistant to the rapid exhaustion normally experienced by proinflammatory macrophages in the TME.

Macrophages play a major role in cancer development and progression. Based on their high degree of plasticity, initial antitumour functions are often subject to rapid protumoral conversion^8,10^. Major efforts in cancer research have tried to reverse this conversion by increasing proinflammatory signalling^1,2,4,9^. In this context, it has been long known that proinflammatory activation, in particular with IFNγ, equips macrophages with tumoricidal activity^3,14^. Yet, efforts to harness this activity for antitumour therapy by increasing IFNγ signalling *in vivo*^17^, stimulating monocytes *ex vivo* with IFNγ^6^, or transducing them with proinflammatory CAR-vectors^12^ have remained clinically disappointing due to the rapid exhaustion of proinflammatory macrophage activity in the tumour^6,7^.

An explanation for this might lie in the nature of proinflammatory macrophage activation itself, which intrinsically leads to the concomitant engagement of immunosuppressive and exhaustion pathways^10,79,80,82^. This includes IFNγ-induced monocyte-derived AHR^+^IL4I1^+^ TAMs that are associated with T cell exhaustion and poor prognosis in many cancers^10,38^. In addition, IFNγ signalling also weakens macrophage stamina by preventing cell cycle progression^19^, favouring cellular aging and compromising metabolic energy efficiency^75,77^. As an indication of this, we show that IFNγ-stimulated WT macrophages acquired IFN-I and senescence signatures that have been associated with cellular aging and protumoral macrophage activity^75–77,79,80^. Interestingly, CAR-activated gene expression^12,81^ overlapped with these IFNγ-induced gene sets, suggesting similar exhaustion mechanisms that might explain poor persistence of CAR-macrophages in patients^7^. Furthermore, IFNγ-driven metabolic reprogramming toward glycolysis is associated with immunosuppressive metabolites^17,20,53,89^, drastically lowers ATP yields and reduces metabolic flexibility to utilize alternative nutrient sources^20,54^. This renders IFNγ-activated macrophages highly glucose-dependent and compromises their metabolic competitiveness against aggressive tumour cells in a nutrient-deprived TME^23,24,90^. Taken together, these mechanisms indicate that proinflammatory IFNγ stimulation comes at the high price of concomitant immunosuppression and cellular exhaustion, presenting an intrinsic dilemma to all therapeutic efforts exploiting proinflammatory activation for antitumoral macrophage activation.

Here we show that activation of stemness in human macrophages can resolve this dilemma by uncoupling the deleterious effects of IFNγ stimulation from proinflammatory antitumour activity. We show that deletion of the self-renewal restricting TFs *MAF* and *MAFB* in human macrophages activates similar self-renewal circuits as previously reported by us in mice^25,26^. Remarkably, we now show that this also resulted in activation of additional stemness features including transcriptional pathways of mitochondrial function and biogenesis that enabled resistance to IFNγ-induced exhaustion. This resonates with recent findings that position mitochondrial health at the core of cellular fitness in the TME^55,56,91–93^. Based on this, mitochondrial transfer has been explored as a therapeutic option^94^, and indeed the transfer of healthy mitochondria could convey stemness features and antitumour efficacy to chronically exhausted T cells^57^. Here we show that stemness activation in human macrophages directly engaged core pathways of mitochondrial biogenesis including *mTORC1*, *YY-1*, *PPARGC1A*, and *NRF1*^62,63^. This resulted in enhanced mitochondrial functionality without the need of mitochondrial transfer, as indicated by MMP, morphology, and respiratory activity. Moreover, enhanced mitochondrial fitness led to reinvigoration of macrophage energy metabolism characterized by increased OXPHOS and ATP generation, despite proinflammatory activation with IFNγ.

Furthermore, we observed activation of cytoprotective signalling pathways in human MAF-DKO macrophages, involving targets of *TP53*, *FOXO* and *NRF2* TFs that mediate cellular longevity by enabling cells to resist redox and replication stress^65–69,72,73^. This might explain the remarkable ability of MAF-DKO macrophages to cope with IFNγ-induced cellular aging mechanisms, high ROS levels and continued proliferation. Surprisingly, NRF2 has been previously reported to protect protumoral TAMs^95^. Our results now suggest that the activation of stemness renders features of cytoprotection available to antitumoral macrophages that were previously thought to be a hallmark of protumoral TAMs.

This remarkable and unique response of stemness-activated human macrophages to IFNγ stimulation thus uncouples activation from concomitant induction of exhaustion. Hence, stemness appears to enable energy efficiency and an extended antitumoral activity in IFNγ-activated macrophages as demonstrated by the rapid, effective, and persisting antitumour activity against several tumour cells *in vitro* and the reduced tumour growth and extended survival of tumour-bearing mice.

These unique traits have major implications for macrophage-based cancer therapies as they address the central problem of exhaustion that has hampered previous trials. This makes the advantages of macrophages such as their ability to infiltrate solid tumours accessible to therapeutic exploitation. Furthermore, the cell-autonomous, CAR-independent antitumour activity extends the utility of CAR-dependent therapies, which are vulnerable to antigen heterogeneity and antigen escape of tumour cells^13^. Consequently, this positions stemness engineering in macrophages as a promising strategy to overcome the major restrictions in maintaining their antitumour activity and to enhance the efficacy of macrophage therapies against solid tumours.

## Supporting information

Extended Video 1 (K562 alone control)

Extended Video 1 (K562 with WT macrophages)

Extended Video 1 (K562 with DKO macrophages)

## Methods

### Cell lines

A549 lung adenocarcinoma cell line was a gift from Mirko Theis, Carl Gustav Carus Uniklinikum Dresden. SK-OV-3 ovarian adenocarcinoma cancer cells and K562 chronic myelogenous leukaemia cells were a gift from Jiri Eitler. A549 and SK-OV-3 cells were transduced with a MOI of 10 with a lentivirus encoding firefly luciferase (fLuc) coupled with a P2A region with a monomeric red fluorescent protein (mRFP) or an Incucyte Nuclight NIR lentivirus driven under an EF-1a promoter (4805, Sartorius). Transgene-expressing cells were derived from sorted single cell clones or by puromycin selection before further use. Cell lines were maintained in complete RPMI (RPMI 1640, 21870076, ThermoFisher), supplemented with 100 units/mL penicillin and 100 µg/mL streptomycin (15140122, ThermoFisher), 2 mM GlutaMAX (35050038, ThermoFisher), and 1 mM sodium pyruvate (11360-039, ThermoFisher).

### Cryopreservation

Harvested EB suspension cells were centrifuged at 1400 rpm for 5 min, and cell pellets were resuspended at 1-10 x 10^6^ cells per cryovial in 1-1.5 ml of Bambanker Serum-Free Cell Culture Freezing Medium (Nippon Genetics, BBH01). Cryovials were stored at -70°C for 24 h before transfer to liquid nitrogen for long-term storage. For cell thawing, cryovials were submerged in a pre-heated water bath (37°C) until only small ice crystals remained. Afterwards, cells were added to fresh RPMI complete medium and centrifuged at 1400 rpm for 5 min.

Cells were resuspended in the final differentiation medium and plated for culture.

### Culture and maintenance of human iPSCs

iBCRT (BIHi001-A-1 (RRID:CVCL_TZ91)) human iPSCs were a gift from Ralf Kühn and cultured under feeder-free conditions in StemFlex Medium (A3349401, ThermoFisher) on vitronectin-coated plates (A14700, ThermoFisher). Cells were passaged with ReLeSR (05872, STEMCELL Technologies) when reaching 60-80% confluency and maintained in the presence of 10 µM ROCK inhibitor Y-27632 (Cay10005583, Biomol) on the day after passaging. Single-cell suspensions of iPSCs were obtained by incubation with Accutase (SF006, Merck).

### Differentiation of human macrophages from iPSCs

Directed differentiation of iPSCs was performed as described previously with modifications^29,30^. In brief, embryoid bodies (EBs) were formed by seeding a single-cell suspension of WT or MAF-DKO iPSCs at a density of 12,500 cells/well into ultra-low attachment U-shaped-bottom 96-well plate (174925, ThermoFisher) resuspended in EB medium consisting of StemFlex Medium supplemented with 50 ng/mL BMP-4 (314-BP, R&D Systems), 20 ng/mL SCF (255-SC, R&D Systems), 50 ng/mL VEGF (AF100-20A, Preprotech), and 10 µM Y-27632 (Cay10005583, Biomol). The 96-well plate was centrifuged for 3 minutes at 100 g to collect the cells at the bottom. These were cultured for four days (37°C, 5% CO_2_). After one and two days, half of the EB medium was replaced with freshly prepared EB medium. After four days, EBs were picked manually using a P200 pipette using cell-saver tips, resuspended in EB differentiation medium consisting of X-VIVO-15 medium (BE02-060F, Lonza) supplemented with 2 mM GlutaMAX (35050038, ThermoFisher), 55 nM β-mercaptoethanol (M6250, Sigma-Aldrich), 100 ng/mL human M-CSF (PHC9504, ThermoFisher), 25 ng/mL human IL-3 (203-IL, R&D Systems). EBs were seeded into ultra-low attachment 6-well tissue-culture plates (174932, ThermoFisher) with 8 EBs/well or 90 mm tissue-culture dishes with 24 EBs/dish (174945, ThermoFisher). Two thirds of the culture medium were exchanged with fresh differentiation medium every five days. Production of monocytes/macrophages from EBs started around day 15 post EB seeding, and suspension cells were harvested, counted, and used for immunophenotyping by flow cytometry and further differentiation or analysis. For large scale productions, we alternatively used Cero vertical rotation incubators with 2 EBs/mL, i.e., 96 EBs/44 mL, in EB differentiation medium containing 25 ng/mL human IL-3 and 100 ng/mL human M-CSF or baculovirus-produced, recombinant human M-CSF of equivalent activity (vector Coofy27-SS-hM-CSF-C-EPEA, Joop VandenHeuvel, protein production service Helmholtz, Centre Braunschweig), and an optimized feeding regime after day 10 with 2/3 of medium change every 2.5 days containing two-fold higher M-CSF concentrations.

### Final differentiation of human macrophages

For further *in vitro* experiments, EB-harvested myeloid cells were replated for final macrophage differentiation into ultra-low attachment tissue-culture plates (174931 or 174932, ThermoFisher) in complete RPMI medium, i.e., RPMI1640 medium supplemented with 10% FBS (A15-101, PAA-GE Healthcare; 21870076, ThermoFisher), 100 units/mL penicillin and 100 µg/mL streptomycin (15140122, ThermoFisher), 2 mM GlutaMAX (35050038, ThermoFisher), 1 mM sodium pyruvate (11360-039, ThermoFisher), 200 ng/mL M-CSF (Joop), and 50 ng/mL GM-CSF (300-03, Peprotech), and cultured at 37°C, 5% CO_2_. Partial medium change was performed every 2-3 days by exchanging 2/3 of the supernatant with fresh medium. Macrophages were considered as fully differentiated after 7-8 days post-plating.

### MAFB and MAF editing

CRISPR editing of iPSCs was performed as described previously^96^ using the CRISPR/Cas9 system to delete the coding sequence of *MAF* (Gene ID: 4094, 373 amino acids) and *MAFB* (Gene ID: 9935, 323 amino acids) in human cells. In brief, BIHi001-A-1 were exposed to sgRNA-expressing plasmids targeting *MAF* and *MAFB* (pU6-(BbsI)sgRNA_CAG-venus-bpA, 86985, Addgene). Two sgRNAs were designed per gene to target a sequence in the 5’ UTR at least 20 nucleotides upstream of the start codon and in the 3’ UTR at least 20 nucleotides downstream of the stop codon of each gene using the CrispRGold programme (https://crisprgold.mdc-berlin.de)^97^. The sgRNAs were expressed from the sgRNA-expression vector pU6-(BbsI)sgRNA_CAG-venus-bpA, which also encodes a fluorescent reporter gene (Venus, derived from YFP) for positive selection of successfully transduced iPSCs. Using Lipofectamine 3000 (L3000001, ThermoFisher), BIHi001-A-1 were transfected with sgRNA expression vectors and pCAG-mTrex2-bpA (86984, Addgene) to enhance indel formation. Control cells (WT) were treated with pU6-(BbsI)sgRNA_CAG-venus-bpA, however, without inserting any protospacer sequence. Transfected BIHi001-A-1 cells were treated with 1 µg/mL doxycycline hyclate (D9891, Sigma) to induce Cas9 expression, and were subsequently sorted for expression of the Venus reporter gene. Sorted cells were seeded at low densities to get single-cell-derived colonies. In the first round of transfection, sgRNAs targeting MAF were used and colonies were screened for full-length deletion of the *MAF* CDS using PCR and Sanger sequencing of the edited locus. Subsequently, *MAF*^-/-^-iPSCs were treated similarly with *MAFB*-targeting sgRNAs. Three *MAF*/*MAFB*-DKO (MAF-DKO)-iPSCs clones were isolated (clone 1, 2, or 3, respectively). For clonal analysis of single-cell-derived colonies, genomic DNA was extracted from iPSCs with QuickExtract (LGCQE09050, Merck). PCR was performed with Herculase II (600675, Agilent). PCR products were gel-purified (28704, Qiagen) and sent for Sanger sequencing by commercial providers.

### Colony assay

Differentiated macrophages (1/microwell) were plated onto AggreWell^TM^800 6-well plates (34825, STEMCELL Technologies), and cultured in STEMdiff^TM^ APEL^TM^ 2 Medium (05270, STEMCELL Technologies). Sequential images of individual macrophages and colonies during expansion were taken using the Keyence BZ-X710 microscope.

### Morphological assessment of macrophage-directed differentiating iPSCs

Cells harvested from various differentiation steps up to final differentiated macrophages were morphologically assessed as described previously with minor modifications^98^. In brief, 3 x 10^4^ cells were cytocentrifuged in 100 µL PBS at 500 *rpm* for five minutes (Cytofuge 8, Germany). Following methanol fixation, microslides (CLS294875X25, Merck) were stained with the Kwik-Diff kit (9990700, Epredia), and images were taken at 10x or 40x magnification with the AxiScanZ.1 slide scanner.

### Macrophage polarization

IFNγ stimulation was performed in complete RPMI medium as indicated above, however, with an increased dose of GM-CSF of 100 ng/mL and the addition of 20 ng/mL IFNγ (AF-300-02, PeproTech) in the absence or presence of M-CSF for 16-24 hours as stated. For tumourinduced reprogramming, medium was removed and replenished with A549-cell-conditioned medium (protocol in the Supplementary Methods) at a 1:1 ratio with complete RPMI medium, supplemented with 50 ng/mL GM-CSF, and 200 ng/mL M-CSF (Joop) for 48 hours (scRNA-Seq and LEGENDplex) or 96 hours (Flow cytometry).

### Western blotting and protein lysates

Protein from WT and MAF-DKO iPSCs was extracted by lysing cells in Laemmli buffer (J60015.AD, ThermoFisher). Lysates were homogenized using QIAshredder (79656, Qiagen), denatured, and loaded onto SDS-polyacrylamide gels for electrophoresis (SDS-PAGE), followed by semi-dry blotting. Blots were blocked with 5% non-fat dry milk (68514-61-4, Biomol) in TBS-T (J77500.K2, ThermoFisher), and incubated with anti-MAFB (HPA005653, Sigma) or anti-MAF antibodies (HPA028289, Sigma), then with a secondary antibody, anti-rabbit IgG-HRP (12-348, Sigma), followed by incubation with ECL^TM^ Prime Western Blot detection reagent (RPN2232, GE Healthcare Amersham^TM^).

### Real-time qPCR

Total RNA was isolated from iPSCs or differentiation stages using the RNeasy Mini Kit (74104, Qiagen), or RNeasy Micro Kit (74004, Qiagen), followed by cDNA synthesis using the High-Capacity Reverse Transcription Kit (4368814, Applied Biosystems). Real-time qPCR was performed using the TB Green Premix Ex Taq (RR420A, Tli RNase H Plus, Takara). PCR products were run on a 2% agarose gel (A42135, Invitrogen).

### Flow cytometry

For flow cytometry, cells were blocked with 1:300 human FcR Blocking Reagent (130-059-901, Miltenyi) for 15 minutes at 4°C, washed in FACS Buffer (PBS with 1% FCS and 2 mM EDTA), and stained with antibodies for 60 minutes in the presence of human FcR Blocking Reagent at 4°C (antibodies indicated in Extended Data Table 2). Cells were recorded on a BD LSR Fortessa, a Cytek® Aurora, or a BD MacsQuant instrument and analysed with FlowJo (v10) or OMIQ (Dotmatics). Cell sorting was performed on a BD Aria II or Aria III. For Annexin V staining (640932, Biolegend), tumour-macrophage co-cultures were stained according to the manufacturer’s instructions. For mitochondrial analysis, isolated cells were stained using 100 nM Mitotracker Deep Red FM (8778S, Cell Signaling), and 200 nM Mitotracker Green FM dyes (9074, Cell Signaling).

### Click-it reactions

Cells were treated with 20 µM OPP (C10458, ThermoFisher) or 10 µM EdU (C10632, ThermoFisher) according to the manufacturer’s instructions. After washing, cell pellets were fixed and permeabilized using the BD Cytofix/Cytoperm^TM^ Fixation/Permeabilization Kit (554714, BD Biosciences) preceding analysis.

### Cytotoxicity assays

For *in vitro* analysis of cytotoxic competence, A549, SK-OV-3, or K-562 cells were transfected with a NIR-expressing plasmid as described before. NIR-expressing tumour cell targets were plated at 1 x 10^4^ cells per 96-well (665090, Greiner), and co-cultured with WT or MAF-DKO macrophages at different ratios. NIR^+^ cells were quantified with an Incucyte SX5 instrument and normalized to NIR^+^ cells at seeding. The medium was changed every 24 hours.

### Cytokine/chemokine analysis using LEGENDplex

Legendplex analysis was used to quantify cytokine concentrations with a customized panel of human cytokines from LEGENDplex (1647, Biolegend) or LEGENDplex Human Free Active/Total TGF-β1 (740449, Biolegend) in undiluted culture supernatants according to the manufacturer’s instructions and acquired on a LSR Fortessa. Cytokine quantification was done using the LEGENDplex^TM^ Data Analysis Software.

### Intracellular ROS assay

Macrophages were plated in a 96-well plate (665090, Greiner) and incubated with 10 µM dihydrorhodamine 123 (DHR-123, D23806, ThermoFisher) in macrophage differentiation medium in a SX5 Incucyte instrument. Green fluorescent intensity based on DHR-123 oxidation was assessed to quantify intracellular ROS production upon addition of 20 ng/mL IFNγ or 1.6 µM phorbol-12-myristate-13-acetate (PMA, P1585, Sigma). To control signal specificity quenching of further ROS production was performed using 10 mM of N-acetyl cysteine (A7250-5G, Merck).

### Seahorse analysis

A Seahorse XF96 Bioanalyser (Agilent) was used to determine OCR. Cells were washed in assay medium (XF DMEM, 103575-100, Agilent) with 10 mM glucose (103577-100, Agilent), 1 mM sodium pyruvate (11360070, Gibco), and 2 mM L-glutamine (25030081, Gibco) at a pH 7.4 at 37°C before being plated onto Seahorse cell culture plates at 1-2 x 10^5^ cells/well. After adherence and equilibration, cellular OCR and ECAR were measured using the Seahorse XF Cell Mito Stress Test Kit (103015-100, Agilent) with addition of 2.5 µM oligomycin, 2 µM carbonyl cyanide 4-(trifluoromethoxy) phenylhydrazone, and 0.5 µM antimycin A or rotenone according to the manufacturer’s instructions.

### NAD^+^ analysis

Macrophages treated for 16 h with 20 ng/ml IFNγ, 50 ng/ml GM-CSF and 200 ng/ml M-CSF were harvested for lysis and NAD+ Extraction according to manufacturer (MAK460, Merck). Fluorescence at 530/585 nm was measured from undiluted samples with a Tecan Plate Reader.

### Caspase 3/7 analysis

*Macrophages treated for* 16 h with 20 ng/ml IFNγ, 50 ng/ml GM-CSF and 200 ng/ml M-CSF, washed and then co-incubated with K562 tumour cells for 48 h at a 10:1 ratio. Media was completely removed and treated for 30 min with 2 μM CellEvent Caspase 3/7 Green Detection Reagent (C10423 ThermoFisher), according to manufacturer’s manual. Caspase 3/7 positive macrophages were discriminated from NIR+ K562 tumour cells by Cell detection software from the Incucyte SX5.

### Mitochondrial analysis

For mitochondrial analyses, macrophages were co-cultured with A549 cells as described before. Co-cultures were established either in µ-Slides (80826, ibidi) for Mitotracker staining and live-cell imaging or in tissue culture-treated 96-well plates for flow cytometry. On day two, macrophages were stained with 200 nM MitoTracker^TM^ Green FM (9082, Cell Signaling Technology). Live-cell imaging was performed using a Dragonfly spinning disk confocal microscope (Oxford Instruments) equipped with a 64x silicone immersion objective. Mitochondrial membrane potential was assessed using the JC-1 assay (551302, BD Biosciences). Cells were incubated with 2 μM JC-1 dye and analysed by flow cytometry on a BD LSR Fortessa. The mitochondrial polarization state was determined by calculating the JC-1 aggregate-to-monomer fluorescence ratio.

### Microscopy image processing

Confocal images were deconvoluted and rendered using Huygens Professional (version 3, Scientific Volume Imaging). For volume reconstruction, movie making and maximum intensity projection images were processed using Vision4D Arivis Software (Version 4.1.2, Zeiss). Rendered images were subjected to an established mitochondrial segmentation pipeline to derive segmented images and mitochondrial features^99^.

### Macrophage functional in vitro assays

Differentiated macrophages were incubated for 1-2 hours with pHrodo Green *E*. *coli* Bioparticles (P35366, ThermoFisher) in phenol-free medium. Phagocytosed bioparticles were measured by flow cytometry. For Cathepsin B activity, macrophages were fixed in BD Cytofix/Cytoperm Buffer (15827868, BD Biosciences) for 20 minutes at 4°C. Then, cells were washed and resuspended in primary anti-Cathepsin B antibody (1:100, 31718S, Cell Signaling) for one hour at 21°C followed by a 30-minute incubation with anti-rabbit-AF488 secondary antibody (A-11008, ThermoFisher). For lysosomal staining, macrophages were stained in 266 ng/mL acridine orange solution (6130, Bio-Rad) for 20 minutes at 37°C. Stained cells were counter-stained with 1 µg/mL Hoechst 33342 (62249, ThermoFisher) for 10 minutes prior to cytocentrifugation and imaged on a Zeiss Inverse Apotome.

### Bulk RNA sequencing

Fragments were aligned to the human reference genome (hg38) with GSNAP (v2020-12-16). Ensembl gene annotation was used to detect splice sites. Uniquely aligned fragments were counted with featureCounts (v2.0.1). Sequence libraries underwent a quality control with RNA-SeQC (v1.1.8). Normalization of the raw fragment counts was done with DESeq2 (v1.24.0) and IHW (v1.12.0) in R (v3.6.3) and based on the library size and testing for differentially expressed genes (DEGs) between the two genotypes (MAF-DKO vs. WT at two hours of stimulation). For GSEA, DEGs were analysed in gseapy (v1.1.5). Leading edge genes were selected for a z-score-based heatmap. The GSEA network was built based on the DEGs and the 200 topmost NES in MAF-DKO macrophages. The Jaccard Index was used for the edges network (v3.5) and Louvain Community Algorithm (v1.0.0b1) for clustering. TF activity was calculated with decoupler (v2.9.7) using three replicates.

### Sample processing and scRNAseq analysis

RNA was isolated and formaldehyde-fixed with the Chromium Fixed RNA Kit (Human Transcriptome, 10x Genomics). For library preparation, samples were thawed and resuspended in a hybridization mix for barcoding. Afterwards, barcoded cells were pooled and subjected to encapsulation. Gel Beads-in-Emulsion (GEMs) were then pre-amplified for eight cycles, and the library amplification was done for nine cycles. Sequencing was done on a NovaSeq 6000 S4 v1.5 (Illumina) with paired-end 100 bp reads to a total of 1.28 x 10^6^ reads. FASTQ files were generated and aligned to the genome with Cell Ranger (v7.1.0). The log1p transformation on minimum values of total counts and gene counts per cell was used as thresholds for filtering cells. The pct_mitochondrial was set at 20% and a doublet score of 0.3 using Scanpy (v1.10.3). Doublet prediction was done with Cell Ranger. Filtered samples were normalized using Scanpy. The integrated dataset was scaled, and a UMAP dimensionality reduction was performed using the top 50 principal components. Leiden clustering was performed with a spatial resolution of 0.4. GSEAs were performed with gseapy (v1.1.5) configurated with the signal-to-noise method. All scores of gene sets and signatures were calculated with score_genes. TF activity was performed using pydecoupler (v1.1.0).

### Mice

NOD/SCID Il2rg-/-hIL3-hGMCSF-hSF (NSG-SGM3 or NSGS) mice were obtained from the Jackson Laboratory. Unless indicated otherwise, 6-20-week-old mice of both sexes were used for experiments and maintained in pathogen-free conditions in individually ventilated cages under a 12 h – 12 h light-dark cycle at a temperature of 21 ± 1°C and a humidity of 40-70%. Mice were fed with autoclaved food and water *ad libitum*. This study complied with all ethical regulations involving experiments with mice, and all experimental procedures performed were approved by the Institutional Animal Care and Use Committee of the Landesdirektion Dresden (TVV6/23). Mice were randomized to allow sex- and age-matching across groups. No animals were excluded from analysis.

### Bioluminescent in vivo imaging

Mice underwent bioluminescent imaging at the indicated time points using D-Luciferin (25836, Cayman Chemical) according to the manufacturer’s instructions. In brief, mice received 3 mg D-Luciferin in 100 µL sterile PBS via intraperitoneal injection in the lower right quadrant using a 30G insulin needle (BD Micro-Fine+). Then, mice were anesthetized using 2% isoflurane (Piramal Critical Care Germany) and the bioluminescence image was acquired 10 minutes after intraperitoneal injection using an AMI HTX Spectral Imager.

### Subcutaneous tumour experiments

A549 fLuc^+^mRFP^+^ (P10-24) were used. Human iPSC-derived macrophages were harvested from the Cero bioreactor at day 20 of production and used together with macrophages harvested at day 15 of production that were further expanded for five days in the absence of EBs (pool day 15+5 with day 20). At collection, macrophages underwent proinflammatory activation at 1-2 x 10^6^ macrophages/mL in 90 mm^2^ ultra-low adherence plates or in the Cero using 400 ng/mL M-CSF (Joop), 100 ng/mL GM-CSF, and 20 ng/mL IFNγ in X-VIVO-15 medium for 16 hours. To remove residual IFNγ, macrophages were washed three times before being co-injected with A549 cells at an E:T ratio of 5:1 for the TME and exhaustion (5 x 10^6^ macrophages with 1 x 10^6^ A549 cells) or at an E:T ratio of 50:1 for the subcutaneous killing experiments (1 x 10^7^ macrophages with 2 x 10^5^ A549 cells) in 150 µL complete X-VIVO-15 medium supplemented with 400 ng/mL M-CSF (Joop) and 50 ng/mL GM-CSF into the right flank. For calliper-based assessment of tumour growth, a modified ellipsoidal formula was used^100^. Mice were sacrificed at the indicated time points or euthanized once termination criteria were reached.

### Multiplex immunohistochemistry

Fresh tissue samples were prepared as described previously^101^. In brief, tumour-resident macrophages were characterized by multiplex immunohistochemistry (mIHC). To assess exhaustion and proliferation of human macrophages, CD11b, SLAMF7, HLA-DR, Ki-67, cleaved Caspase 3, and DAPI were used. Detailed information on staining specifications is provided in Extended Data Table 3. Staining of formalin-fixed, paraffin-embedded (FFPE) tissue sections was performed as described previously^101,102^. The protocol concluded with nuclear counterstaining using DAPI (Sigma Aldrich), mounting with Fluoromount-G® (SouthernBiotech), and storage in the dark at 4°C. Whole slide scanning was performed using the Phenollmager HT 2.0 Imaging System (Akoya Biosciences) at 100x magnification. Regions of interest (ROIs) were defined using Phenochart^TM^ Software (Akoya Biosciences). Multispectral images (MSIs) were acquired at 200x magnification. Spectral unmixing and image generation was done as described previously^103^. A Python script (v3.13) was written for automated image analysis of the CD11b, HLA-DR, and SLAMF7 area quantification. Images were applied in grayscale onto the tumour-macrophage mask, and a Gaussian blur and multi-Otsu thresholding was used to create a binary mask for each marker. The mask was applied to focus the analysis on the tumour area for which Napari was used for manual definition based on the DAPI signal. The area was calculated to correspond to the percentage of the positive area for each marker. Multi-Otsu thresholding was used as follows: HLA-DR = thresholds [-2], CD11b/SLAMF7 = thresholds [-3]. Packages used: mamba (v2.2.1), napari (v0.6.1), numpy (v2.2.6), scikit-image (v0.25.0), scipy (v1.15.2), pandas (v1.5.3), tqdm (v4.67.1), seaborn (v0.13.2), opencv (v4.11.0), pyqt (v5.15.11), openpyxl (v3.1.5), scikit-learn (v1.2.2), and statannotation (v0.5.4). For Ki-67 scoring or assessment of cleaved Caspase 3 (Cas3), i.e., quantification of Ki-67^+^CD11b^+^ or Cas3^+^CD11b^+^ macrophages, the area calculation tool within QuPath (v0.6.0) was used to calculate the tumour area within each image based on DAPI signal intensity. Afterwards, the Ki-67 or Cas3 signal was counted within each CD11b^+^ macrophage. The final count was normalized to 1 x 10^6^ µm^2^ of analysed tumour area.

### Statistical analysis

All data were tested for normality using a Shapiro-Wilk test and data is presented as median ± IQR or mean ± SD unless indicated otherwise. Statistical analyses were performed using Prism10 (GraphPad). Experiments were reproduced in at least two independent runs with at least three biological replicates. No statistical method was used to predetermine sample size. Sample size was determined by previous experience with similar experimental models in our laboratory^104,105^. Analyses performed include a paired or unpaired Student’s t-test to compare two datasets, one-way ANOVA with Tukey’s multiple-comparison tests for multiple group comparisons. To compare groups over time, the Kruskal-Wallis test and mixed-effect analysis (RELM model) were used. To compare more than two groups, two-way analysis of variance (ANOVA) was used.

### Data availability

All data from this study are available in the article, extended data and supplementary data. All data from RNA sequencing experiments can be accessed from the Gene Expression Omnibus under accessions available upon acceptance. Source data are provided with this paper.

## Acknowledgements

We thank members of the M.H.S. laboratory, Martin Wermke and Martin Bornhäuser for helpful discussion and Vasileia Ismini Alexaki for technical advice on metabolic assays. We thank A. Gompf and K. Bernhardt for flow cytometry, the light microscopy facility (both CMCB Technology Platform, TU Dresden) and the CRTD Animal facilities for mouse husbandry. This work was supported by Institutional grants from TU Dresden and grants from the European Research Council (ERC) (grant agreement numbers 695093 MacAge and 101069255 OncoMac to M.H.S.) and the European Innovation Council (EIC) (grant agreement number 101291635 Stemphage) under the European Union’s Horizon 2020 and Horizon Europe research and innovation program, the Saxocell cluster4future (FKZ 03ZU1111HA), GOBio Initial (FKZ 16LW0263K) and GOBio feasibility (FKZ 16LW0042) grants from the BMBF, and the ForTra gGmbH of the Else Kröner Fresenius Stiftung 2024_EKTP14 (EKFS). The research was further partly funded by the Federal Ministry of Research, Technology and Space and co-funded by the European Commission (01KT2304B to M.S.). In addition, the research was supported by the Federal Ministry of Research, Technology and Space (03ZU2111AB to M.S). J.S. was funded by the German Research Foundation (Clinician Scientist position, SU 1360/1-1). J.S. and S.S. are participants of the national Translational Tandem Programme for Gene- and Cell-based Therapies (nTTP-GCT) – coordinated by the Biomedical Innovation Academy of the Berlin Institute of Health at Charité (BIH) and funded by the Federal Ministry of Education and Research (BMBF) (01BIH_nTTP-GCT_25-07). The funders of the study had no role in the study design, data collection, data analysis, data interpretation or writing of the manuscript. M.H.S. has been funded as an Alexander von Humboldt Professor at the TU Dresden by the Alexander-von-Humboldt Foundation (FRA/1188926). Open Access funding enabled and organized by Projekt DEAL. Illustrations in Extended Data Fig. 1 d,g and 7a were created using <u>BioRender.com</u> under a paid subscription.

## Author contributions

J.N., C.J.B., J.S., and M.H.S. were involved in all aspects of this study, including planning of experiments and investigation, analysis and interpretation of data. J.S. performed the majority of in vivo and J.N. the majority of in culture experiments, J.F., A.S., A.Sc., M.G. and S.Su. assisted with experimental design and performed experiments. M.H. performed the majority of bioinformatic analyses. F.R. and L.E. contributed analyses and discussion of bioinformatic data, respectively. M.F. produced lentiviruses and transduced A549 and SK-OV-3 cells. J.B. analysed microscopy data and provided technical expertise on spectral flow cytometry. Q.L. and S.Sa. provided critical resources. S.Sa. and S.Su. provided supervision and coordination for specific aspects of the study and contributed expertise in key technologies, including animal experimentation. D.I. performed mitochondrial and Seahorse assays. A.St. and M.S. provided critical reagents and performed multiplex immunohistochemistry. All authors provided critical scientific feedback on the results and manuscript. J.N., C.J.B., J.S. contributed to writing the manuscript, J.N and J.S generated figures. M.H.S. conceived the project, supervised the study and wrote the manuscript.

## Additional information

### Supplementary information

The online version contains supplementary material that will be made available upon acceptance.

### Correspondence and requests for materials

These should be addressed to Michael H. Sieweke.

## Extended Data Figure Legends

**Extended Data Figure 1.**
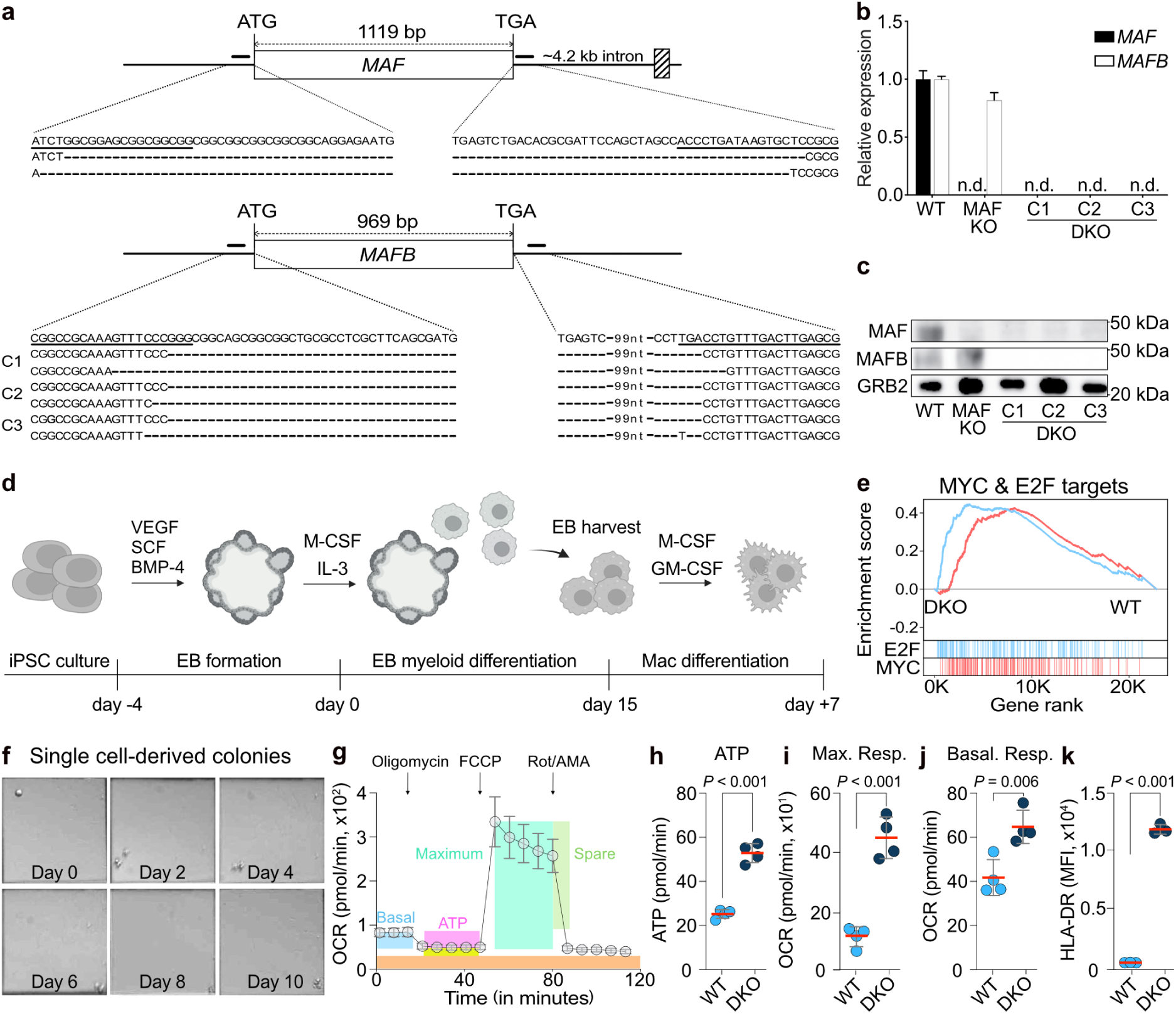
Related to Figure 1. Genetic deletion of *MAF*/*MAFB* activates self-renewal and additional stemness functions. **a,** Gene structure of *MAF* and *MAFB* indicating position of CRISPR/Cas9 target sites in the 5’ and 3’ UTR of both genes, and indels obtained. The first line of each sequence block refers to the wild-type locus, subsequent lines indicate the indels generated by CRISPR/Cas9 editing. Start and stop codons are bold, genomic target sequences are underlined. The protospacer positions are shown as a black line close to the start or stop codon. **b,** gene expression of *MAF* (black) or *MAFB* (white) by qPCR. **c,** Protein expression of MAF and MAFB by immunoblotting. **d,** Schematic of human iPSC-derived macrophage differentiation indicating timing and cytokines **e,** GSEA of *MYC* and *E2F* targets. **f,** Phase-contrast images from single cell cultures of EB harvest-derived WT macrophages at indicated time points. **g,** Schematic of an oxygen consumption rate (OCR) profile from a Seahorse MitoStress test assay. Indicated are readout parameters including basal respiration (Basal), ATP production (ATP), maximum respiration (Maximum), spare respiratory capacity (Spare), and non-mitochondrial respiration (orange), as well as administration of oligomycin, Carbonyl cyanide-4-(trifluoromethoxy)-phenylhydrazone (FCCP), and rotenone/Antimycin A (Rot/AMA). **h-j,** Quantification of ATP production (**h**), maximum respiration (**i**), and basal respiration (**j**) from Seahorse analysis of unstimulated MAF-DKO or WT macrophages. **k,** Flow cytometric quantification of HLA-DR surface expression by MFI. Statistical analysis was performed using unpaired two-tailed t-tests. Data shown as mean ± s.d. Representative results from one of at least three independent experiments. Full gene set descriptions shown in Extended Data Table 4. EB, embryoid body.

**Extended Data Figure 2.**
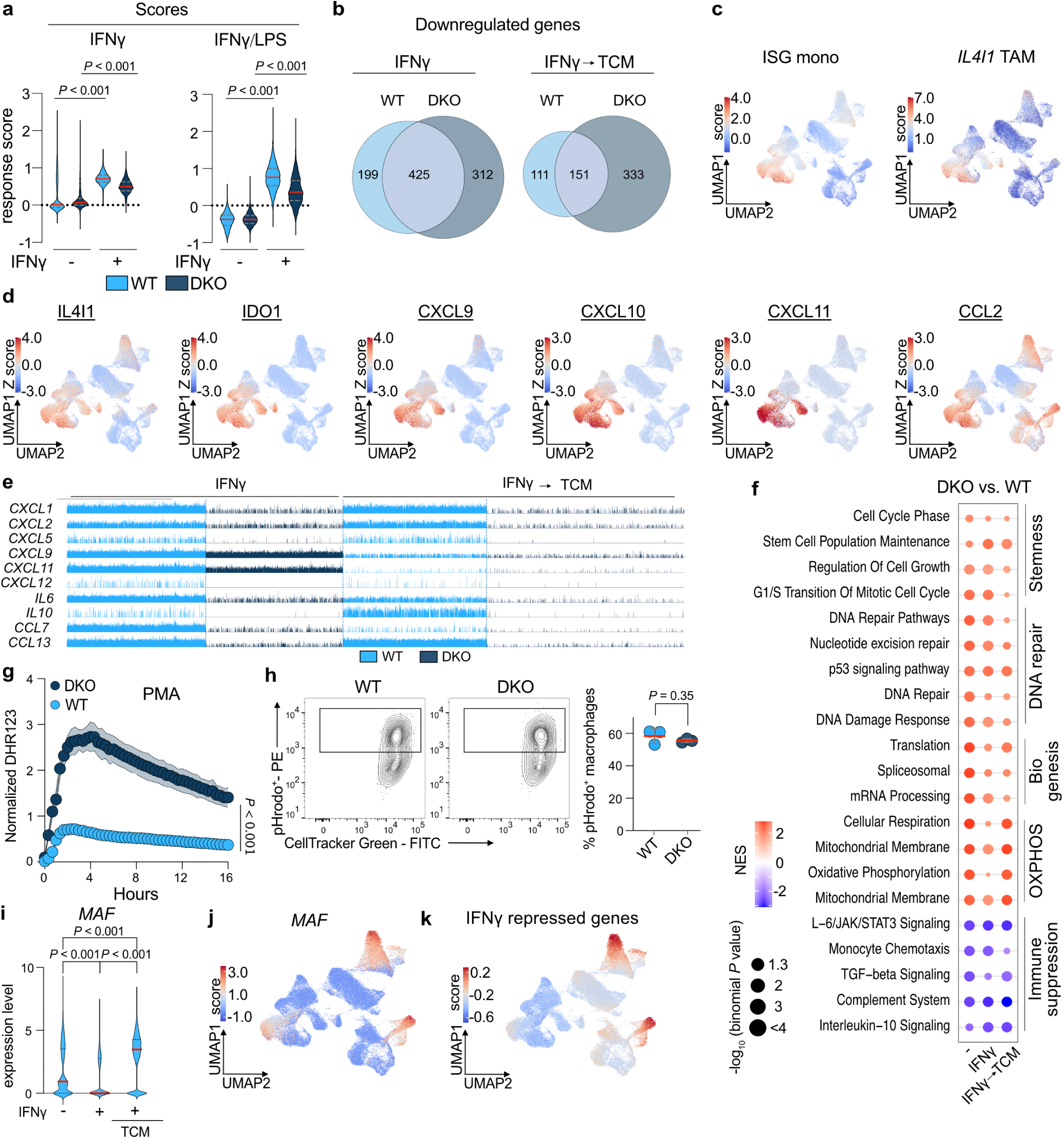
Related to Figure 2. MAF-DKO macrophages show a unique response to IFNγ. **a,** Violin plots showing scores for gene signature of IFNγ^35^ or IFNγ/LPS stimulation^34^ responses in blood-derived human monocytes. **b,** Proportional Venn diagrams for differential IFNγ-downregulated genes of WT and MAF-DKO macrophages in response to IFNγ or IFNγ with subsequent exposure to TCM. **c,** UMAP projection of ISG monocytes (ISG mono) and *IL4I1*^+^ TAMs (*IL4I1* TAM) score^10^. **d,** UMAP projections for indicated genes. **e,** Track plots showing single cell expression of indicated genes in WT and MAF-DKO macrophages upon IFNγ treatment or subsequent TCM exposure. **f,** Bubble plot of gene sets enriched in MAF-DKO over WT macrophages. **g,** Quantification of intracellular reactive oxygen species (ROS) levels in PMA-stimulated WT and MAF-DKO macrophages monitored by DHR fluorescence signal in Incucyte SX5 live-cell imaging at indicated times, expressed as relative green fluorescent units per well normalised to cell confluence, *n* = 3 per group. **h,** Representative flow cytometry plots and quantification of percent pHrodo^+^ of CellTracker Green-labelled macrophages 24 hours after co-culture with pHrodo-labelled A549 cells, *n* = 3 per group. **i,** Violin plot of *MAF* expression in WT macrophages at indicated conditions. **j,** UMAP projection of *MAF* and *MAF*-dependent IFNγ-repressed genes^35^. Statistical analysis was performed using a two-way ANOVA with Tukey’s post hoc t-test (**a,i**), a Mixed-effect analysis (RELM model) to compare two groups over time (**g**), or unpaired two-tailed t-tests (**h**). Data are shown as mean ± s.d. Representative results from one of at least three independent experiments. Full gene set descriptions shown in Extended Data Table 4. TCM, tumour cell conditioned medium. PMA, Phorbol-12-myristat-13-acetat. DHR, dihydrorhodamine 123.

**Extended Data Figure 3.**
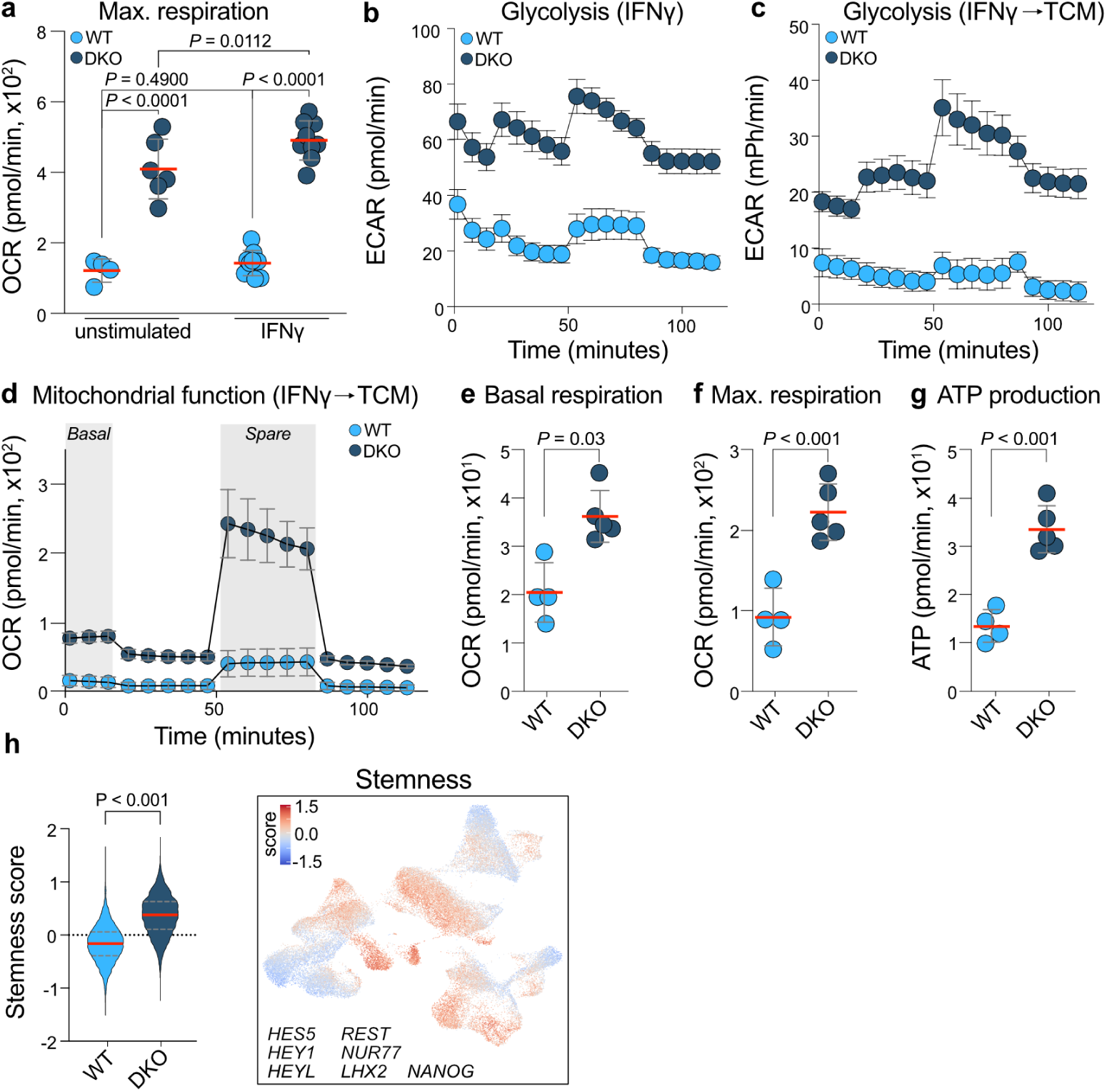
Related to Figure 3. Metabolic characterisation of MAF-DKO macrophages in response to IFNγ and subsequent exposure to TCM. **a,b,** Quantification of maximum respiration and extracellular acidification rate (ECAR) after IFNγ-activation. **c-g,** Quantification of ECAR, OCR, basal respiration, maximum respiration and ATP production in WT and MAF-DKO macrophages treated with IFNγ and afterwards TCM, *n* = 4-6 per group. **h,** Violin plot and UMAP projection of stemness TFs activity score in WT and MAF-DKO macrophages. Statistical analysis was performed using unpaired twotailed t-tests (**e,f,g,h**) or a two-way ANOVA with Tukey’s post hoc t-test (**a**). Data are shown as mean ± s.d. Representative results from one of at least three independent experiments are shown. ECAR = extracellular acidification rate. OCR = oxygen consumption rate. TCM = tumour cell medium.

**Extended Data Figure 4.**
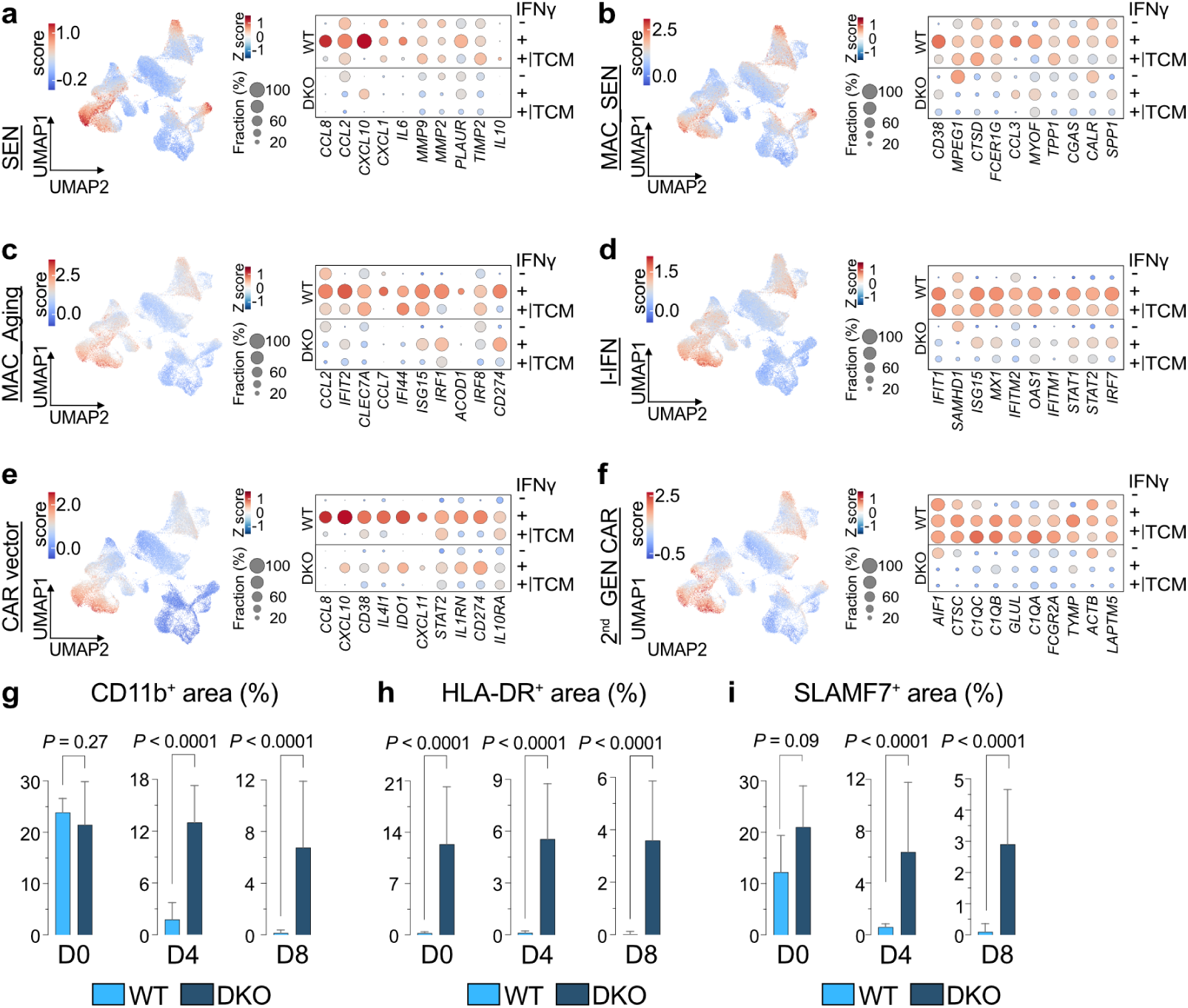
Related to Figure 4. MAF-DKO macrophages resist exhaustion signatures. **a-f,** UMAP projection of published senescence, aging, and activation signatures and bubble plots selected leading edge genes for senescence (SEN) (**a**)^82^, senescent macrophages (MAC_SEN) (**b**)^79^, aging in macrophages (MAC_Aging) (**c**)^75^, type I interferon signatures (IFN-I) (**d**), CAR vector-transduced macrophages (CAR vector) (**e**)^12^, and transcriptional signatures of macrophages transduced with a 2^nd^ generation CAR vector (2^nd^ GEN CAR) (**f**)^81^ in WT and MAF-DKO macrophages without (-) or with IFNγ (+) and subsequent exposure to TCM. **g,** Quantification of CD11b^+^, HLA-DR^+^ and SLAMF7^+^ area in subcutaneous A549 tumours at indicated time points after co-transplantation of IFNγ-activated WT or MAF-DKO macrophages with A549 cells, from Figure 4 p-r, representative of two independent experiments, *n* = 18-41 ROIs in at least two mice per genotype and timepoint. Full gene set descriptions shown in Extended Data Table 4. Statistical analysis was performed using unpaired two-tailed t-tests. Data shown as median ± IQR.

**Extended Data Figure 5.**
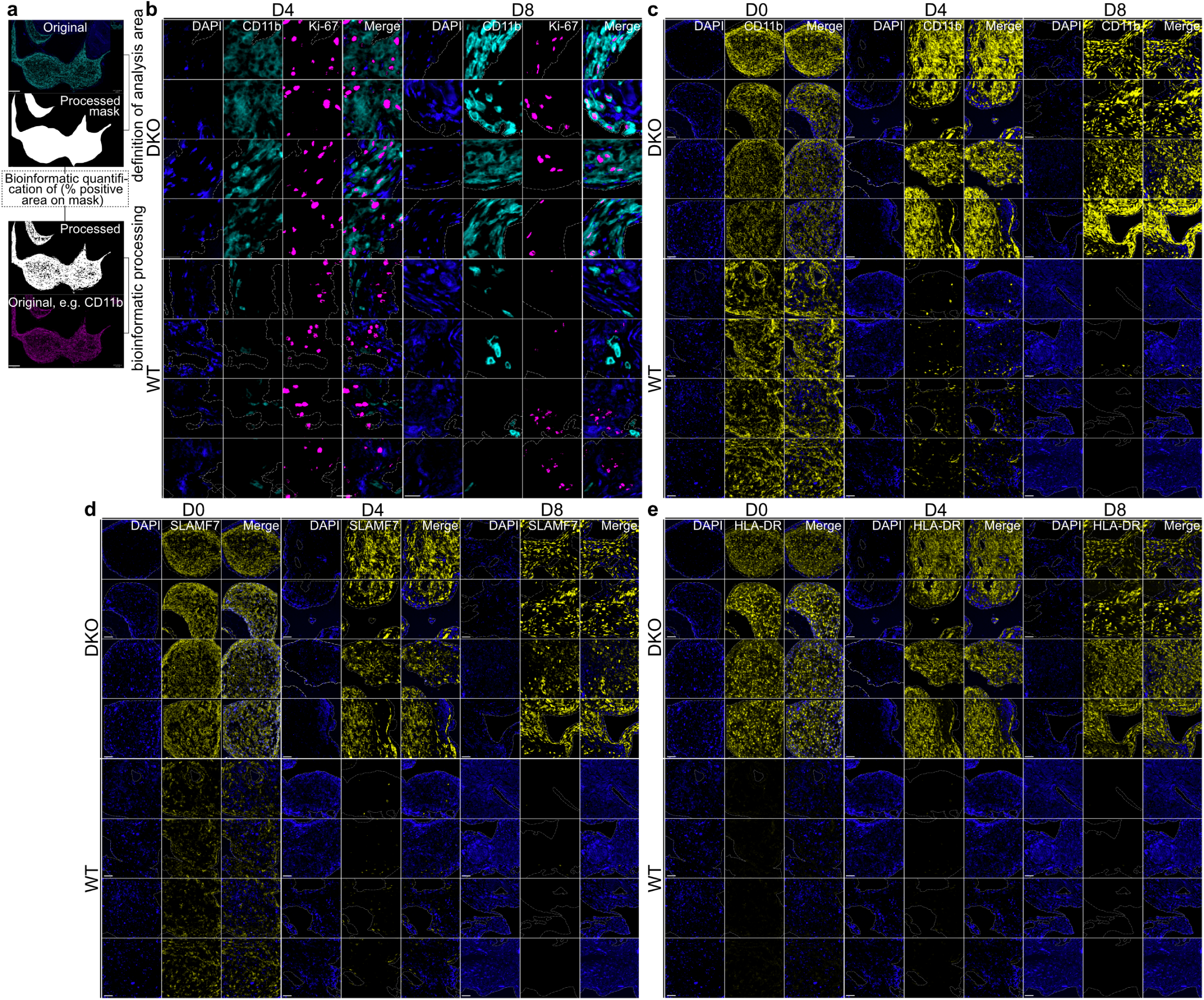
Related to Figure 4. Characterisation of MAF-DKO macrophages *in vivo* and analysis workflow. **a,** Workflow for bioinformatic quantification of fluorescent signals within defined regions of interest (ROIs). A mask was generated from the original microscopy image to quantify the percentage of positive area for specific markers (Ki-67, CD11b, SLAMF7, and HLA-DR). **b-e,** extensive overview of Ki-67^+^CD11b^+^ macrophages (**b**), CD11b^+^ macrophages (**c**), SLAMF7^+^ macrophages (**d**), HLA-DR^+^ macrophages (**e**).

**Extended Data Figure 6.**
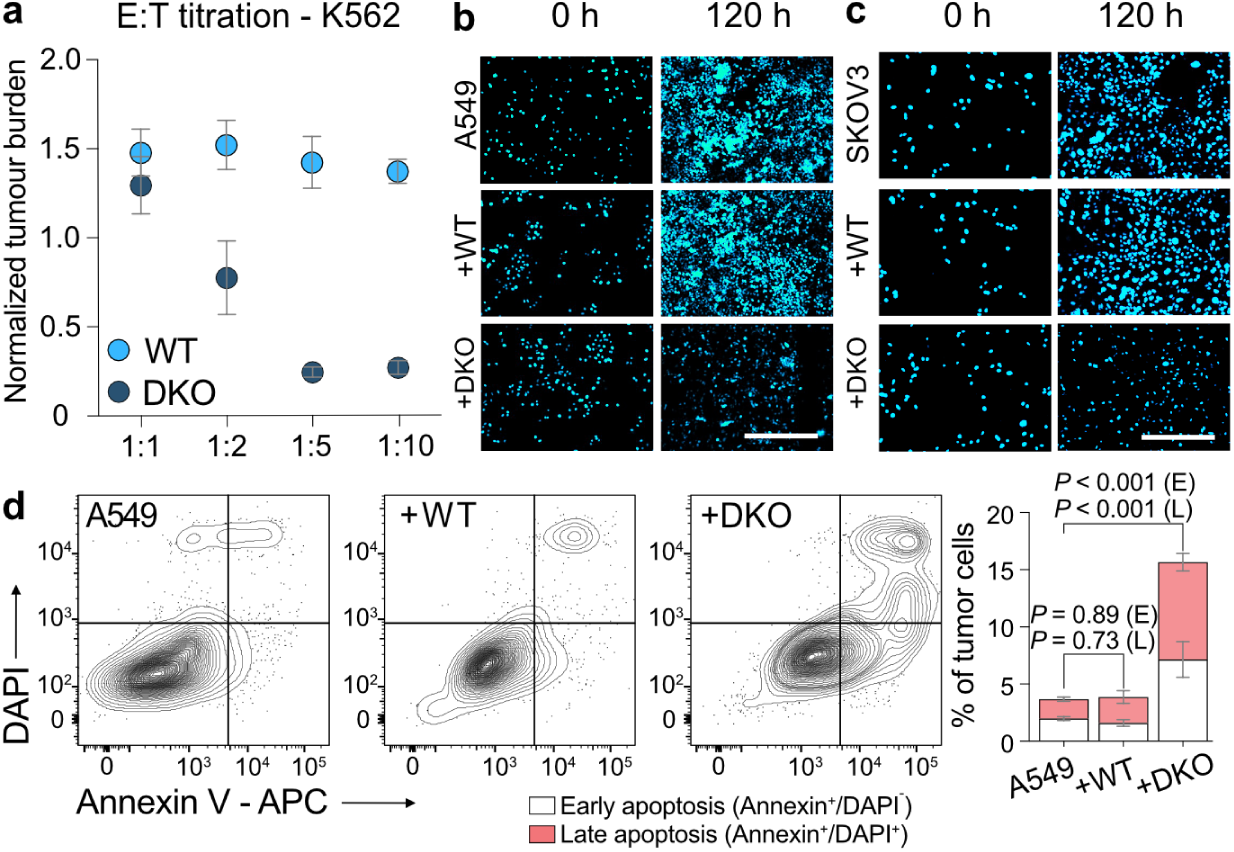
Related to Figure 5. Antitumour function and activation persistence of MAF-DKO macrophages. **a,** Effector-to-target ratio (E:T) titration of Incucyte-based cytotoxicity assay quantifying NIR^+^ K562 target cells normalised to time = 0h after 24 hours of co-culture with WT or MAF-DKO macrophages, *n* = 3 per group. **b,c,** Representative images of A549 and SKOV-3 monocultures and co-cultures with WT and MAF-DKO macrophages at seeding (0 hours) and endpoint (120 hours). Scale bar = 20 μM. **d,** Representative FACS plots and quantification of Annexin V and DAPI signal in CD45^-^CD11b^-^ A549 cells after five days of monoculture (left) or co-culture with WT (middle) or MAF-DKO macrophages (right) at an E:T of 10:1. Quantification (far right) represents the percentage of early or late apoptotic tumour cells, *n* = 3 per group. Statistical analysis was performed using a two-way ANOVA with Tukey’s post hoc ttest. Data are shown as mean ± s.d. and are representative of at least three independent experiments.

**Extended Data Figure 7.**
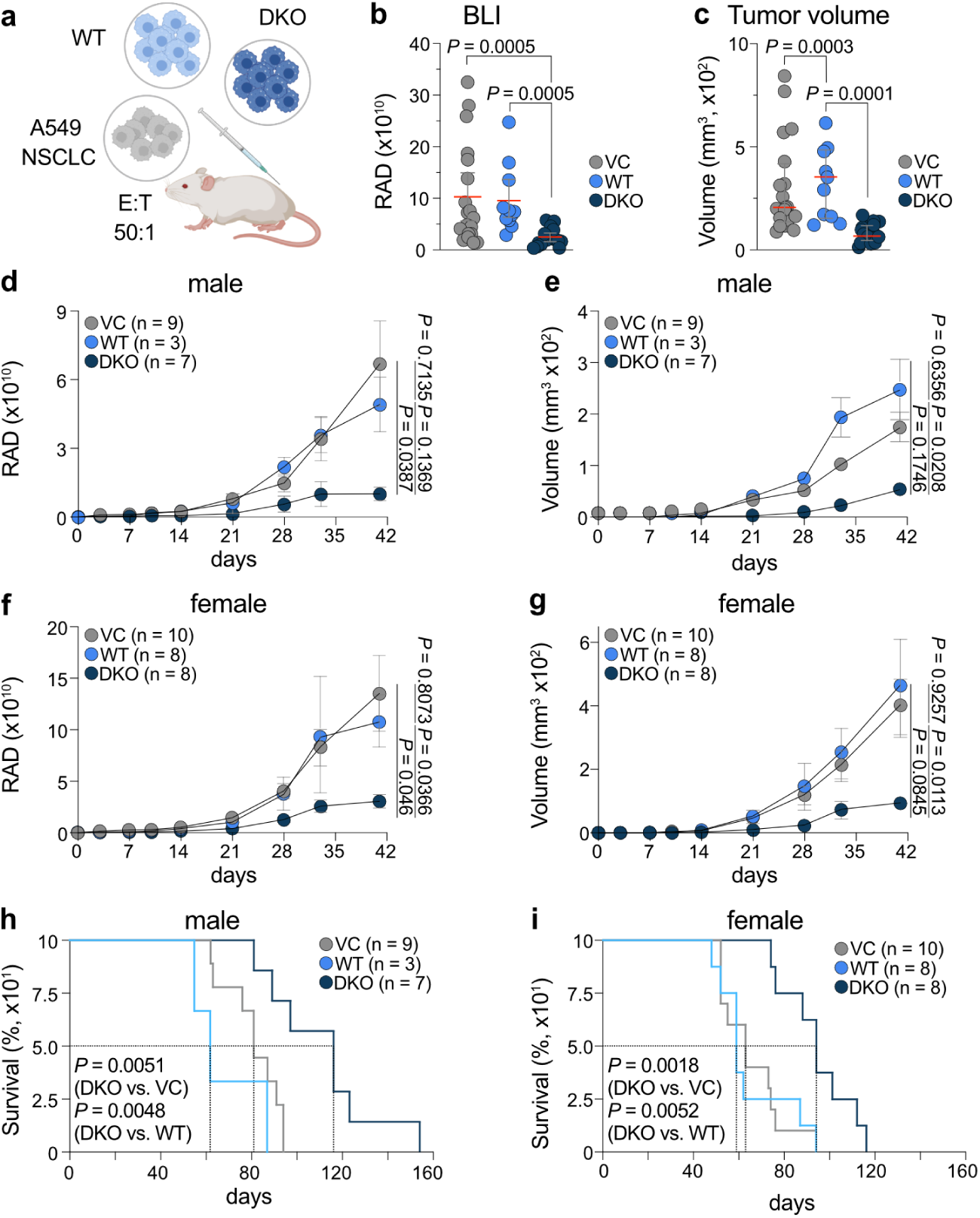
Related to Figure 5. MAF-DKO macrophages exert persisting antitumour activity *in vivo* against NSCLC irrespective of sex. **a,** Experimental schedule of subcutaneous co-injection of A549 cells either with vehicle or WT/MAF-DKO macrophages at an E:T ratio of 50:1. Volumetric assessment^100^ or bioluminescent imaging (BLI) was performed at the indicated time points and mice were euthanized once termination criteria were met. **b,c,** Analysis of BLI (flux p/s) (**b**) or tumour volume (mm^3^) at day 41 (**c**) of all mice irrespective of sex and treatment with vehicle (*n* = 19), WT (*n* = 11), or MAF-DKO macrophages (*n* = 15). **d,e,h,** Male-specific analysis of average BLI (flux p/s) (**d**) or volume in mm^3^ (**e**) or survival (**h**) of mice treated with vehicle (*n* = 9), WT (*n* = 3), or MAF-DKO macrophages (*n* = 7) over time. **f,g,i,** Same as (**d,e,h**) for female-specific analysis of mice treated with vehicle (*n* = 10), WT (*n* = 8), or MAF-DKO macrophages (*n* = 8) over time. Data are median ± IQR (**b**-**g**) and were analysed by Kruskal-Wallis test (**b,c**), a two-way analysis of variance (ANOVA) with Tukey’s post hoc t-test (**d-g**), or log-rank Mantel-Cox test (**h,i**).

**Extended Data Table 2:**
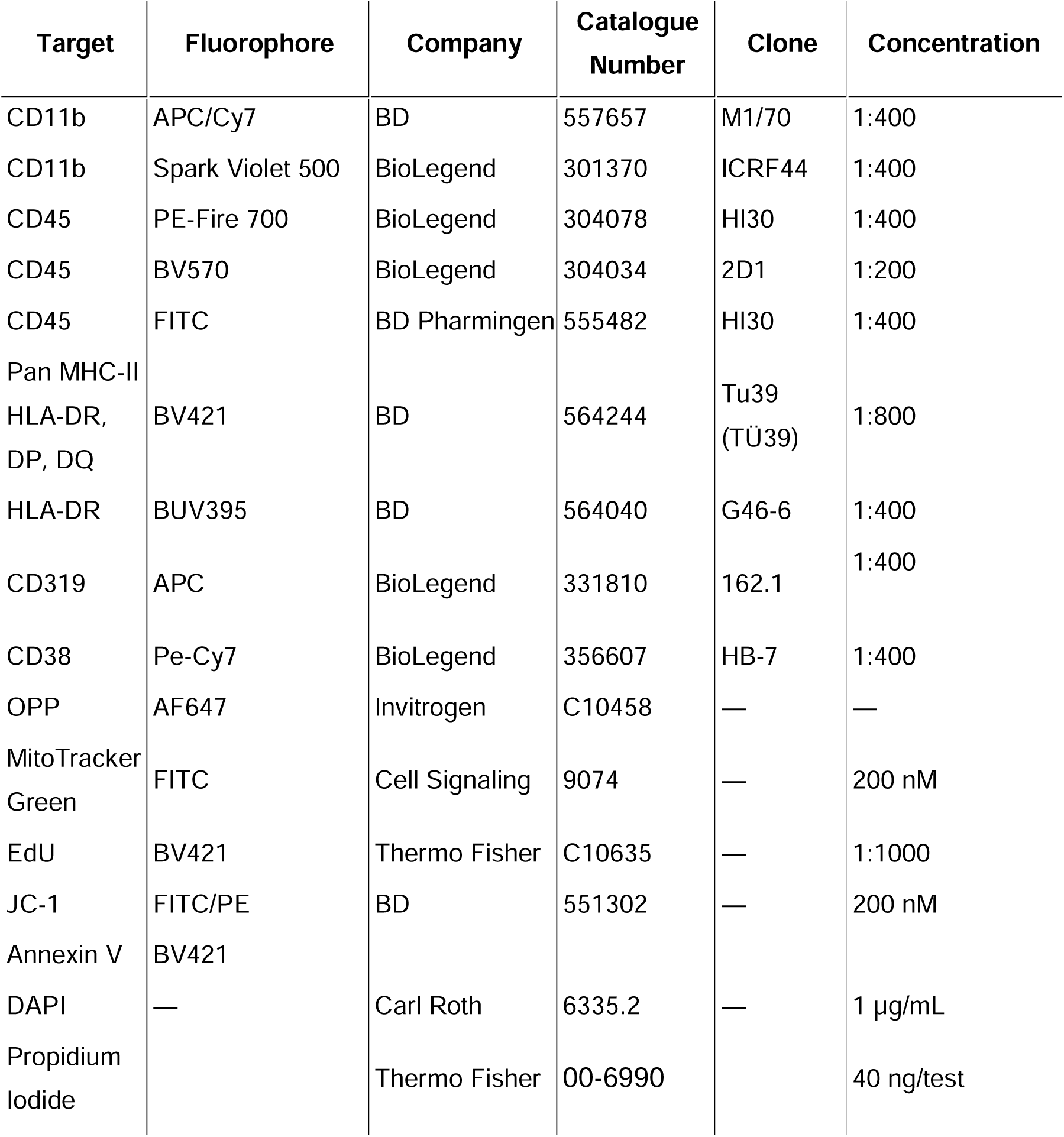
Flow cytometry Antibodies.

**Extended Data Table 3:** Immunofluorescence Antibodies.

| Antibody | Primary Antibody Dilution | Incubation Temperature & Time | Fluorophore | Fluorophore Dilution | Clone | Supplier |
| --- | --- | --- | --- | --- | --- | --- |
| CD11b | 1:150 | 36 °C, 32 min | 650 | 1:100 | D6X1N | Cell Signaling Technology, Inc. |
| Cleaved Caspase-3 | 1:150 | 36 °C, 32 min | 540 | 1:150 | 5A1E | Cell Signaling Technology, Inc. |
| HLA-DR | 1:75 | 36 °C, 32 min | 620 | 1:100 | E9R2Q | Cell Signaling Technology, Inc. |
| Ki-67 | 1:1000 | RT, 16 min | 570 | 1:1000 | D3B5 | Cell Signaling Technology, Inc. |
| SLAMF-7 | 1:100 | 36 °C, 32 min | 690 | 1:200 | E5C4M | Ventana Medical Systems, Inc. |

**Extended Data Table 4: Full gene sets used in this manuscript (separate document)**

