## Supplementary figures and images for "Stemness overcomes IFNγ-induced antitumour exhaustion in human macrophages"

### Extended Data Table 1

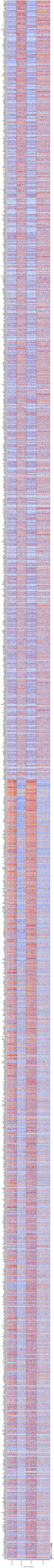
